# A new structural paradigm for outer membrane protein biogenesis in the Bacteroidota

**DOI:** 10.1101/2025.02.17.638638

**Authors:** Xiaolong Liu, Luis Orenday Tapia, Justin C. Deme, Susan M. Lea, Ben C. Berks

## Abstract

In Gram-negative bacteria the outer membrane (OM) is the first line of defence against antimicrobial agents and immunological attacks. A key part of OM biogenesis is insertion of OM proteins (OMPs) by the β-barrel–assembly machinery (BAM). Here we report the cryo-electron microscopy (cryoEM) structure of a BAM complex isolated from *Flavobacterium johnsoniae*, a member of the phylum Bacteroidota that includes key human commensals and major anaerobic pathogens. This BAM complex is radically different from the canonical *Escherichia coli* system and includes an extensive extracellular canopy that overhangs the substrate folding site and a subunit that inserts into the BAM pore. We find that two of the novel subunits involved in forming the extracellular canopy are essential for BAM function. For one of these subunits isolation of a suppressor mutation allows the separation of their essential and non-essential functions. The need for a highly remodelled and enhanced BAM complex reflects the unusually complex membrane proteins found in the OM of the Bacteroidota.

## Main text

Gram-negative bacteria are distinguished by the presence of an OM at the cell periphery^1^. This membrane is the site at which the bacterium interacts with its environment (or host if a pathogen), and provides the first line of defence against antibiotics, mechanical stresses, and immunological attacks. These functions depend on the presence of proteins that either form a transmembrane β-barrel (Outer Membrane Proteins, or OMPs) or are anchored to the membrane by a lipid tail (lipoproteins). Sophisticated pathways are required to target, insert, and fold these proteins. Although these pathways are well-characterised in *E. coli* and related organisms^2–4^, the mechanistic details of OM protein biogenesis in other major Gram-negative phyla remain unclear. Knowledge of OM biogenesis is critically important in combating antimicrobial resistance through identifying new antimicrobial targets (e.g. ^5,6^) or through compromising the OM barrier function.

The BAM complex is central to OM biogenesis since it catalyses the folding and insertion of nascent OMPs into the OM. The core of the BAM complex is BamA. In *E. coli* the BAM complex also contains four periplasmic lipoprotein subunits, BamB, BamC, BamD, and BamE^7^. Only BamA and BamD are essential for BAM function and the roles of the remaining subunits remain poorly defined^8^. BamA is a member of the Omp85 family of 16-stranded OMPs^8^ and is related to the central subunit of the machinery that inserts β-barrel proteins into the mitochondrial OM^9^. The BamA barrel has a periplasmic extension composed of five POTRA (<u>po</u>lypeptide <u>tr</u>ansport–<u>a</u>ssociated) domains to which the lipoprotein subunits bind^10–12^. Within the BamA barrel the seam between the first and last strands is unusually short and weak and can open^10,12^ allowing the exposed N-terminal strand of the BamA barrel to pair with the C-terminal strand of an incoming substrate OMP^13,14^. This structure in turn templates insertion and folding of successive strands of the nascent OMP through β-augmentation. The result is the formation of a hybrid barrel between BamA and the client OMP that is only resolved when the OMP barrel has been completed and closes to release it from BamA^15,16^.

It is currently assumed that the canonical BAM system of *E. coli* (BAM_Ec_) is representative of the structural organisation and mechanism of BAM complexes across the bacterial domain. However, the OM proteins found in some bacterial phyla exhibit considerably greater structural diversity than the *E. coli* OM proteome, raising the possibility that the capabilities of the BAM machinery in these phyla might be augmented relative to BAM_Ec_. One obvious case where this might apply is the phylum Bacteroidota (formerly Bacteroidetes). The Bacteroidota are abundant Gram-negative commensals in the human gut and other human microbiomes^17^ and include major human opportunistic anaerobic pathogens responsible for sepsis (e.g. *Prevotella* species, *Bacteroides fragilis*) and severe dental disease (*Porphyromonas gingivalis* and *Tannerella forsythia*). In the Bacteroidota many OMPs are characterised by large extracellular regions that are far more substantial than those found on *E. coli* OMPs^18–20^ and so the Bacteroidota BAM system must be capable of handling the biogenesis of such proteins. In addition, and also in contrast to *E. coli*, the Bacteroidota possess abundant cell surface lipoproteins (SLPs) which the BAM complex is considered a prime candidate to export^18,21^. Importantly, both of these biosynthetic requirements are required to assemble the SUS nutrient uptake systems that are a characteristic and highly abundant feature of the Bacteroidota OM, because these are centred on a complex between a SLP (SusD) and an OMP with large extracellular regions (SusC) ^18,22^. A further intriguing aspect of BAM in the Bacteroidota is a possible connection with a Bacteroidota-specific protein transport system called the Type 9 Secretion System (T9SS) ^23^. In those Bacteroidota possessing the T9SS, two of the essential T9SS components are encoded at the *bamA* locus^24^ suggesting a functional link between Bacteroidota BAM and Type 9 protein export.

To investigate the nature of the Bacteroidota BAM system we have isolated and characterized the BAM complex from the T9SS-possessing bacterium *Flavobacterium johnsoniae*.

## Structure of the *F. johnsoniae* BAM complex

We isolated the native *F. johnsoniae* BAM complex (BAM_Fj_) using an affinity tag fused to the BamA (Fjoh_1690) protein. Biochemical (Fig.1a) and structural (Fig. 1b Extended Data Figs. 1, 2, and 3a, and Table S1) analysis revealed that the BAM_Fj_ complex contains five proteins in addition to BamA. One of these proteins could be assigned as BamD (Fjoh_3469). However, the remaining co-purifying proteins were unrelated to known BAM subunits from other organisms, nor did they include the anticipated T9SS components. We name the novel BAM_Fj_ subunits BamF (Fjoh_1412), BamG (Fjoh_0823), BamM (‘Metal ion-containing’, Fjoh_0050), and BamP (‘Periplasmic’, Fjoh_1771). Smaller BamA-containing complexes that appear to be fragmentation products of the BAM_Fj_ complexes were also present in the sample (Extended Data Fig. 1).

**Fig. 1.**
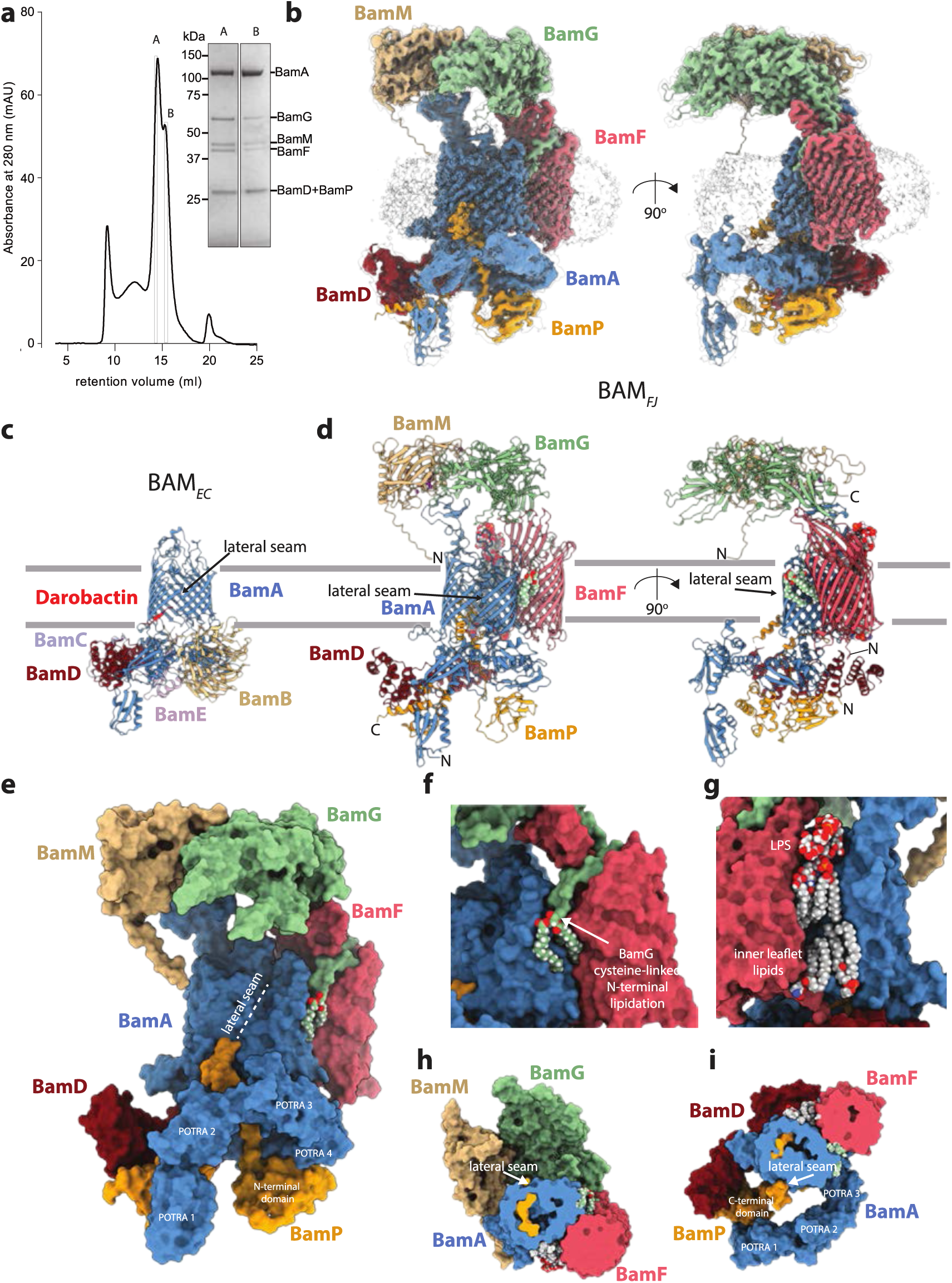
| Structure of the *F. johnsoniae* BAM complex. **a**, Size exclusion chromatography profile of the purified BAM_Fj_ complex and Coomassie-stained SDS–PAGE gel of the indicated fractions. Band identities were assigned on the basis of peptide fingerprinting. Fraction A was used to determine the structure of the full BAMFj complex and fraction B to determine the structure of the BamAP complex. Similar data were obtained for three independent preparations. **b**, CryoEM volume for the BAM_FJ_ complex. The volume is shown at a high contour level (coloured) and at a low contour level (semi-transparent grey with AlphaFold-modelled protein structures placed in the density in cartoon representation). **c,d** Comparison of (**c**) the most similar *E. coli* BAM complex structure (darobactin-bound complex, PDB 8adi) with (**d**) the *F. johnsoniae* BAM complex. Structures are shown in a cartoon representation with lipids and metal ions in spacefill atom representation coloured by element, and Darobactin-9 bound to the *E. coli* structure in red. The inferred position of the OM is indicated by grey lines. **e-i**, Views of the *F. johnsoniae* BAM complex with protein components in space filling representation coloured by subunit and lipids shown as atom spheres coloured by element. **e**, View of the whole complex looking towards the lateral seam. The complex is oriented as in (**c**) Centre. **f,g**, Ordered lipid molecules on opposite sides of the BamA-BamF interface. **f**, The N-acyl and S-diacylglyceryl groups attached to the N-terminal cysteine of the lipoprotein BamG. **g**, The resolved portion of a lipopolysaccharide (LPS) molecule in the outer leaflet of the OM and two phospholipid molecules on the inner leaflet of the OM. **h**, View from the periplasm towards the exterior. The periplasmic side of the complex has been cut away as far as the midpoint of the membrane. **i**, View from the exterior towards the periplasm. The extracellular side of the complex has been cut away as far as the midpoint of the membrane.

As in the *E. coli* BAM complex, BamA forms the core of BAM_Fj_ to which the other subunits are directly or indirectly attached (Fig. 1c). However, whilst the accessory subunits of the *E. coli* complex are all located in the periplasm (Fig. 1c), BAM_Fj_ has a remarkably different organisation in which only BamD and BamP are periplasmic or part periplasmic proteins (Fig. 1b,d,e). Uniquely, the BamF subunit is a transmembrane OMP, whilst BamG and BamM are SLPs that together form an extensive extracellular structure. BamF is bound to the ‘rear’ of the BamA barrel relative to the lateral seam. The interaction between BamA and BamF is reinforced by lipid binding on either side of the subunit interface. On one side these interactions are provided by the phospholipid tail of BamG (Fig. 1f) and on the other by an ordered lipopolysaccharide molecule in the outer leaflet of the membrane and two ordered phospholipid molecules in the inner leaflet (Fig. 1g). BamG and BamM interact with each other to form a long canopy-like structure on the extracellular side of the OM that extends from the ‘rear’ of BamA across the BamA barrel and out beyond the lateral seam to cover the position in the membrane where client OMPs assemble on BamA (Fig. 1b,d,e,h). The canopy is positioned at an approximately constant height of 40 Å above the inferred position of the membrane bilayer and delineates a ca. 3,000 Å^3^ space above the membrane surface. The canopy is anchored to the BAM_Fj_ complex through binding to extracellular ‘pillars’ provided by the BamA and BamF subunits. At the periplasmic side of the membrane the folded domains of the novel BamP subunit are bound to BamD and to the POTRA domains of BamA but elaborate a loop that enters the interior of the BamA barrel (Fig. 1b,d,e,h,i). The more membrane distal POTRA 1-3 domains of BamA, together with the C-terminal portion of BamP, are poorly resolved in the structure and are modelled in all figures by placement of AlphaFold^25^ structures into the EM map (Fig. 1b).

In structurally characterised BAM complexes the lateral seam of BamA has been observed to be either open or closed. In our BAM_Fj_ structure BamA is in the closed state. The lateral seam is held together by two inter-strand hydrogen bonds (Fig. 1d,e) as in the darobactin-bound state of BAM_Ec_^5^ which is its closest structural homologue (Fig. 1c).

The interstrand loops of the *F. johnsoniae* BamA barrel exhibit several notable structural changes relative to the canonical *E. coli* protein (Fig. 2a,b). First, loop 2-3 in is enlarged by 15 residues relative to the *E. coli* protein, forming a short amphipathic structure along the periplasmic face of the OM that extends away from the BamA barrel. Second, loop 9-10 is highly elongated (84 residues in BamA_Fj_ relative to 8 residues in BamA_Ec_) and extends out into the extracellular space where it folds into a β-sheet domain that provides the binding platform for BamM. Finally, as in other BamA proteins, loop 11-12 enters the barrel lumen where it contacts the barrel wall through a conserved ‘VRGF/Y motif’^11,26,27^(the actual sequence being _779_LRGY_782_ in BamA_Fj_). However, in BamA_Fj_ this loop is extended within the barrel pore by 16 residues relative to BamA_Ec_, forming an additional strand loop that extends across and fills the extracellular end of the pore. Notably this additional loop contacts the most deeply inserted piece of BamP. AlphaFold 3 modelling^28^ indicates that all three of these loop structures are highly conserved across Bacteroidota BamA proteins, although only the proteins from Flavobacteria include the BamM-binding domain at the tip of loop 9-10.

**Fig. 2.**
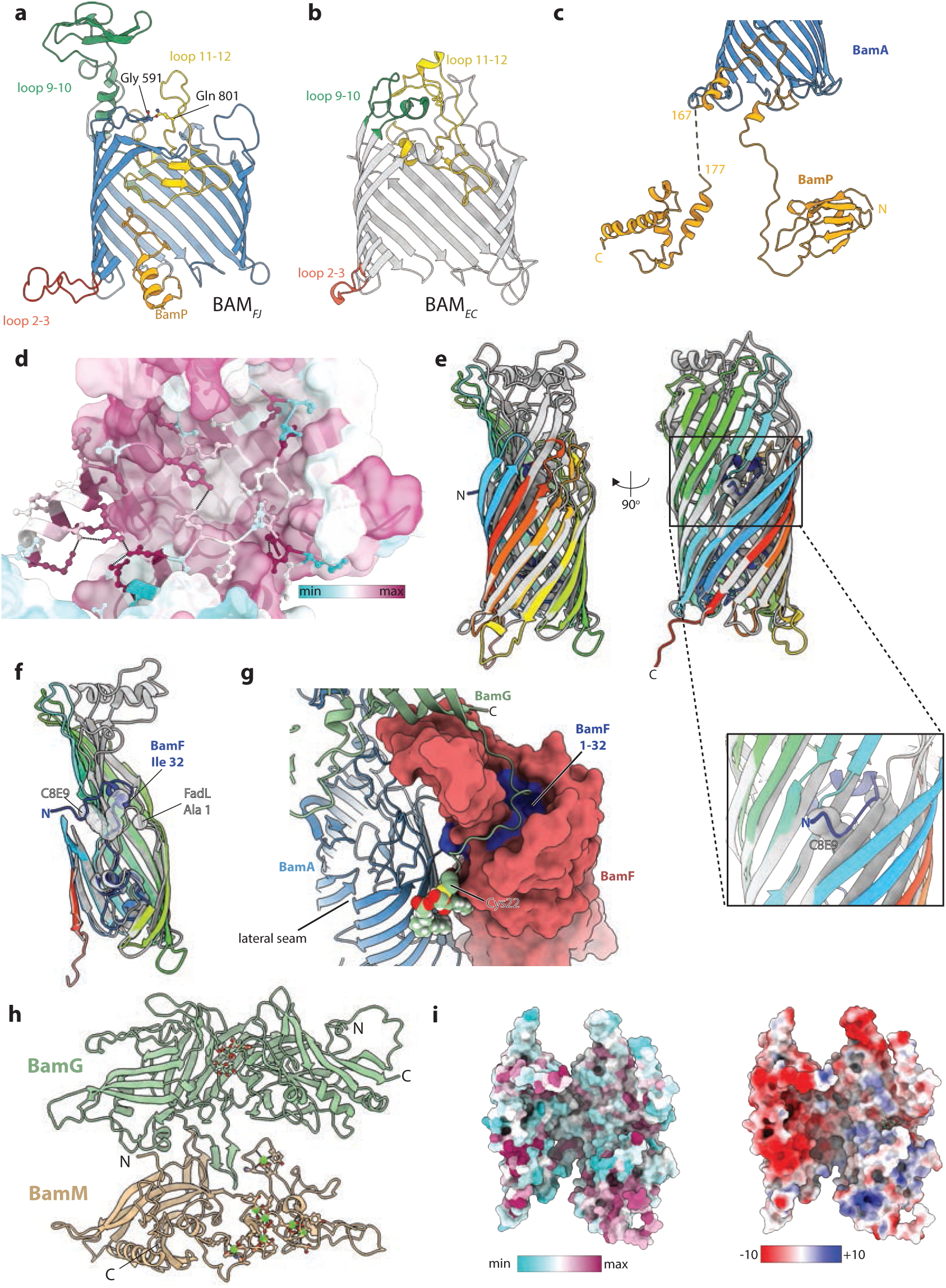
| Structural features of the *F. johnsoniae* BAM complex subunits. **a,b**, Comparison of the *F. johnsoniae* and *E. coli* BamA barrels. The barrels are shown in the same orientation in cartoon representation with the strands closest to the viewer removed and loops highlighted as annotated. **a**, *F. johnsoniae* BamA together with the loop of BamP within the BamA barrel. The residue Gln801 that is substituted in the *bamG* suppressor mutant, together with residue Gly591 to which the side chain of Gldn801 makes a main chain hydrogen bonding interaction, are shown in ball and stick representation and coloured by element. **b**, *E. coli* BamA (PDB: 8adi). **c**, Carton representation of BamP (yellow). The BamA barrel (blue) is shown in cutaway for orientation. **d**, Sequence conservation and intra-chain interactions of the inter-domain loop of BamP (cartoon with ball and stick side chains) within the BamA barrel (grey surface). **e,f** Superimposition in cartoon representation of BamF (chainbows colouring) and FadL (grey, PDB 3dwo). A proposed substrate-mimicking C_8_E_9_ detergent molecule in FadL is shown in grey spheres. In (**f**) the front walls of the barrels, oriented as in (**e**) Left, are cut away and the N-terminal amino acid of FadL together with the equivalent sequence position residue in BamF are shown as spheres. **g**, View from outside the cell showing how the N-terminal region of BamG is bound by BamF. BamF is shown in surface representation with the N-tail (residues 1-32) coloured blue. Partial structures of BamA and BamG are shown in cartoon representation with the N-terminal cysteine of BamG and its attached lipid groups shown as atomic spheres and coloured by atom. **h**, The BAM_FJ_ extracellular canopy in cartoon representation viewed from BamA with the BamF-proximal end at the bottom of the panel. Glycosylation of BamF and bound calcium ions and their co-ordinating residue side chains in BamM are shown in ball and stick representation. **i**, Surface conservation (Left) and electrostatics (Right) of the extracellular canopy in the same orientation as (**h**).

The novel BamP subunit has a tripartite structure in which the N-terminal and C-terminal regions form small structured domains that are linked by an extended central region (Fig. 2c). The most N-terminal part of the protein folds into a carboxypeptidase-like regulatory domain structure (a 4+3 β-sandwich with Greek-key topology) that binds to POTRAs 4 and 5 of BamA (Figs. 1d,e and 2c). From this domain the central region extends up into the BamA barrel, which it penetrates as far as loop 11-12 (Fig. 2a) while making conserved contacts with the interior of the barrel (Fig. 2d). The loop then exits the open periplasmic end of the lateral seam and runs back into the periplasm (Fig. 1d,e and 2a,c). BamP ends in a three-helix C-terminal domain that is sandwiched between, and thus links, BamA POTRA 1 and BamD (Fig. 1d,e,i). This blocks the direct contact between POTRA 1 and BamD that occurs in BAM_EC_. In the position seen in our structure BamP would block binding of the incoming substrate OMP at the lateral gate. It would also block binding of the BAM-specific antibiotic darobactin^5^(Fig. 1c) potentially explaining the insensitivity of Bacteroidota to this antibiotic^5,29^.

BamF is a member of the FadL family of 14-stranded OMPs. These proteins are characterised by a lateral opening in the transmembrane barrel and a long N-terminal tail that threads through the barrel pore to reach the extracellular side of the membrane^30^ (Fig. 2e,f). Canonical FadL proteins function as transporters for hydrophobic molecules. In these proteins the lateral opening acts as a conduit to move hydrophobic substrate molecules from the protein interior into the membrane bilayer^31^. However, in BamF the N-terminal tail is extended and now threads through the lateral opening with the N-terminal residue of the tail touching BamA (Fig. 2e-g). Many additional contacts between BamF and BamA are present and span the entire width of the bilayer. BamF also makes limited contact with BamD through the final three amino acids of its C-tail (Fig. 1d Right). BamF is *O*-glycosylated on the periplasmic end of the N-tail.

The extracellular portions of BamF function to anchor BamG to the BAM_Fj_ complex through extensive contacts. Strands 3 to 7 of the BamF barrel extend into the extracellular space to form the pillar onto which the proximal folded end of BamG docks (Figs. 2e-g). The lipidated N-terminal tail of BamG is an extended piece of polypeptide that runs around the pillar in a deep groove in the BamF surface, before exiting towards BamA (Fig. 2g) in order to position the three acyl chains to pack between the BamA and BamF barrels (Figs. 1g and 2g).

BamG is an elongated molecule with a complex fold that resembles chondroitin sulfate-binding carbohydrate binding domain at the BamF-proximal end (Fig. 2h and Extended Data Fig. 3b). BamG is *O*-glycosylated on the side facing BamA (Fig. 2h).

BamM is composed of two domains (Fig. 2h). The N-terminal domain has a peptidyl-prolyl isomerase (PPI)-like fold (Extended Data Fig. 3c). The C-terminal domain adopts a novel fold that contains no well-defined secondary structural elements but which is structured in part by the presence of seven metal ions (Fig. 2h and Extended Data Fig. 3c). Based on their co-ordination chemistry we assign these metal ions as calcium ions. The phospholipid tail of BamM is not resolved. Nevertheless, the N-terminus of the polypeptide can be fully traced and is appropriately positioned to allow the attached lipid groups to insert in the OM (Fig. 1b,d,e). BamG and BamM are arranged side by side along their long axes and are in contact for about half their length around their centres (Fig. 2h). A large part of the interaction interface between the two proteins arises through each subunit attaching a protruding β-hairpin to the other protein (Fig. 2h). The membrane-proximal side of the BamGM unit would be expected to face substrate proteins and has a deep central valley (Fig. 2i). This surface is hydrophilic, highly acidic in the BamM portion, and shows little amino acid conservation (Fig. 2i) suggesting that it does not make highly specific interactions with substrates.

*F. johnsoniae* possesses homologues of BamF, BamG, and BamP (Fig. 3a). The BamF and BamG homologues, hereafter BamF2 and BamG2, are co-transcribed (*fjoh_1686-1685* locus). Notably BamG2 would be unlikely to interact with BamM as it lacks the protruding β-hairpin that BamG uses for this purpose. The *bamP* operon encodes two further BamP homologues, which we term BamP2 (Fjoh_1769) and BamP3 (Fjoh_1770). A further BamP homologue, BamP4 (Fjoh_2401), is coded at another locus. The BamP homologues have related folded domains but markedly diverge in the interdomain loop. With the exception of BamP4, none of these BAM_Fj_ subunit homologues is expressed at an appreciable level in cells cultured on rich medium^32^.

**Fig. 3.**
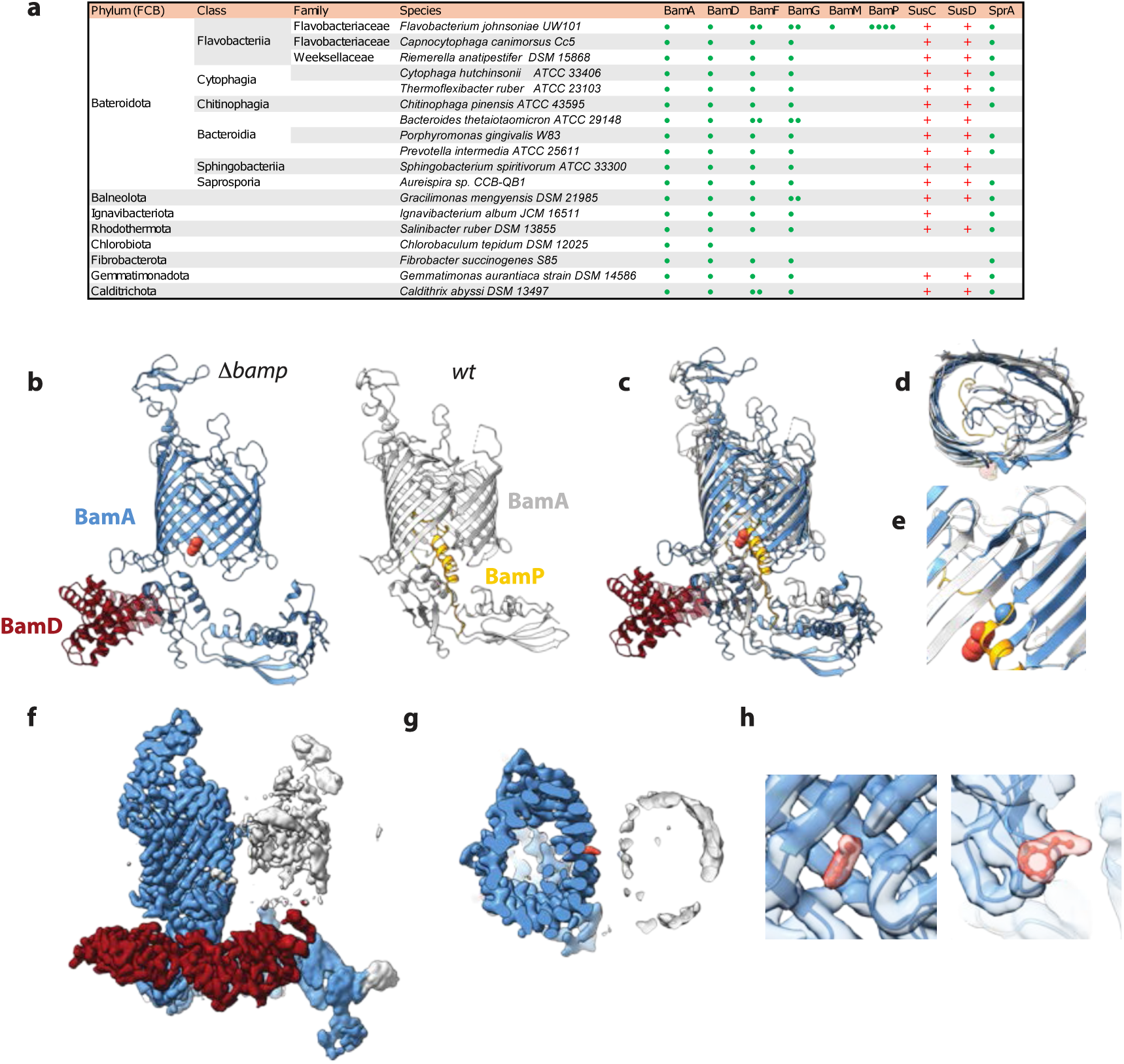
| Phylogenetic distribution of BAM_Fj_ subunits and structural consequences of the loss of BamP. **a**, Distribution of BAM_Fj_ subunit homologues within representative members of the Bacteriodota and wider FCB superphylum. The presence of each copy of a coding gene is indicated by a green dot. Also shown is the distribution across these strains of members of the SusC and SusD protein families (red +) and the 36-stranded T9SS translocon barrel SprA. See also Table S2. **b**, Comparison of the structure of the BamAD complex isolated from a BamP-deleted (Δ*bamP*) background (Left) and a BamAP complex purified from the wt background (Right). The proposed phenylalanine residue is shown in orange spacefill. **c-e**, Overlay of the structures shown in (**b**) aligned on the N-terminal 100 residues of the BamA barrel. **d**, The BamA barrel viewed from the extracellular side of the membrane. **e**, Detail showing the enlargement of the sheet between the BamA barrel N– and C-termini in the absence of BamP and the incompatible binding modes of BamP and the putative phenylalanine ligand (orange spacefill). The C-termini of the BamA models are shown as spheres. **f-h**, Views of the cryoEM volume for the BamAD complex isolated from a BamP-deleted background reveal a partially occupied second barrel incorporated in the micelle. **f**, Side view coloured as in (**b** Left) with unassigned density coloured silver **g**, Top-down view of a slab cut at the level of the putative phenylalanine (orange) bound in the lateral seam. **h**, Two views of the putative phenyalanine bound in the lateral seam and viewed (Left) from the BamA exterior or (Right) perpendicular to the phenylalanine ring and with the BamA barrel cut away to reveal the phenylalanine density.

## BamFG are conserved across the FCB superphylum

BamA, BamD, BamF, and BamG are universally conserved across the Bacteroidota suggesting that they comprise the core components of the Bacteroidota BAM system (Fig. 3a and Table S2). Notably the novel BamF and BamG subunits are also conserved across six of the seven phyla that together with the Bacteroidota comprise the wider FCB (Fibrobacterota-Chlorobiota-Bacteroidota) superphylum (Fig. 3a) indicating that these phyla also possess a Bacteroidota-like BAM complex. Although additional homologues of BamF and BamG are found in *F. johnsoniae* this is not a general feature of FCB BAM systems (Fig. 3a).

Homologues of the full length BamM protein are only found in the genus Flavobacterium, and this is also true of the structures in BamA (elaborated L9-10) and BamG (protruding β-hairpin) that bind this subunit. Thus, BamM is a Flavobacterium-specific accessory subunit. The degree to which BamP is conserved is more difficult to assess because it is composed of a mix of common and poorly defined protein folds. Nevertheless, obvious homologues are restricted to the Family Flavobacteriaceae suggesting that this subunit may not be a general feature of Bacteroidota BAM complexes.

## BamM and BamP are not essential for core BAM_Fj_ function

In order to probe the functional roles of the novel BAM_Fj_ subunits we attempted to disrupt their coding genes. The genes encoding BamM and BamP were successfully deleted either alone or in combination (Extended Data Fig. 4a,b). However, we were unable to delete the genes for BamF or BamG (or in control experiments BamA and BamD) suggesting that these proteins are essential (Extended Data Fig. 4b).

Strains lacking either BamM or BamP or both exhibited no growth defect in rich medium (Extended Data Fig. 4c). They also showed no defect in the canonical BAM function of OMP insertion as assessed through testing for loss of OM integrity by sensitivity to detergents, to EDTA, or to the normally OM-impermeable antibiotic vancomycin^7^(Extended Data Fig. 4d). We raised the possibility above that the Bacteroidota BAM complex could have a role in the biogenesis of SusCD systems or in the operation of the T9SS. However, we observed no inhibition of growth of the deletion mutants on carbon sources (galactomannan and xyloglucan) that require SusCD systems to metabolise^33^(Extended Data Fig. 4e) or of the ability of cells to sustain the T9SS-dependent process of gliding motility^34^ (Extended Data Fig. 4f). Thus, under laboratory conditions BamM and BamP do not detectably contribute to BAM_Fj_ function.

The genes coding for the BAM_Fj_ subunit homologues could also be deleted (Extended Data Fig. 4b). The resulting strains again show no defects in cell growth or OM integrity with the exception that loss of BamP4 results in a modest reduction in sensitivity to EDTA (Extended Data Fig 4c-e).

## Structural consequences of removing the BamP subunit

The central loop domain of BamP is bound at the lateral seam of BamA in a way that would sterically impede hybrid barrel formation with the substrate protein. Consequently, this loop must become displaced during the BAM_Fj_ catalytic cycle. In an attempt to mimic the loop-displaced state we deleted the BamP subunit and structurally characterised the resulting BamA complex.

Although the BamA preparation from the BamP-deleted strain contained all the remaining BAM_Fj_ subunits, only BamAD complexes could be identified by EM following cryo-freezing (Fig. 3b, Extended Data Fig. 5, and Table S1). Fortunately, this loss of the BamFGM subunits does not in itself affect the conformation of the BamA barrel since the barrel conformer does not change between the full BAM_Fj_ complex and a BamAP sub-complex present in the original BAM_Fj_ preparation (Figs. 1a and 3b,c, Extended Data Fig. 6, and Table S1).

In the absence of BamP the BamA barrel remains in the Closed state (Fig. 3b). Indeed, removing BamP allows the C-terminal strand of the barrel to slide along the N-terminal strand towards the periplasm to form an additional hydrogen bond at the lateral seam (Fig. 3e). This change in strand register at the lateral seam results in a limited distortion of the barrel cross-section (Fig. 3d). These structural changes suggest that BamP does not function to lock BamA in the closed state but rather assists in the transition to the open state by partially destabilising the lateral seam.

Unexpectedly the BamAD complex structure contains partial density for a second β-barrel positioned on the lateral seam side of BamA (Fig. 3f,g) as well as unconnected density at the periplasmic side of the BamA lateral seam that we model as a phenylalanine side chain (Fig. 3b,c,e,g,h). The second barrel could correspond either to a second poorly ordered copy of BamA or the averaged density of the mixture of OMPs that proteomics detects as co-purifying with the BamAD complex. It is unclear whether the second barrel and phenylalanine are adventitiously bound or correspond to substrate analogues.

## Assessing the effects of depleting essential BAM_Fj_ subunits

To gain insight into the roles of the essential BamF and BamG proteins we developed a genetic system to allow gene depletion in *F. johnsoniae*. In this system a duplicate copy of the gene of interest is expressed ectopically on the chromosome under the control of a TetR-repressible promoter (Extended Data Fig. 3a-d). Provided expression of this second copy of the gene is maintained by the inclusion of the inducer anhydrotetracyline (aTC) in the growth medium, the native copy of the gene can be deleted. If aTC is then omitted from the growth medium of the resulting strain, the target protein is no longer synthesised and becomes depleted as the cells grow and divide. Using this depletion strategy we confirmed that BamF, BamG and, as a comparator, BamA are essential for growth under standard laboratory conditions (Fig. 4a). In all three cases full depletion of the target protein is apparent by 6 hours after removal of the inducer (Fig. 4b) at which point cell growth slows (Fig. 4a). Within a further two hours the cells become noticeably misshapen (Extended Data Fig. 7e) and start to lose periplasmic contents (Fig. 4b, SkpA lanes). More detailed analysis of the depleted cells by transmission electron microscopy shows that all three depletion strains exhibit a similar perturbed morphology in which the OM no longer buds OM vesicles^35^ but has become deformed by massive blebbing, while the inner membrane remains intact (Fig. 4c). Thus, depletion of any of the three essential BAM_Fj_ subunits leads to gross defects in OM biogenesis. Similar morphological defects in the OM have been reported in *E. coli* following BamA depletion^36^.

**Fig. 4.**
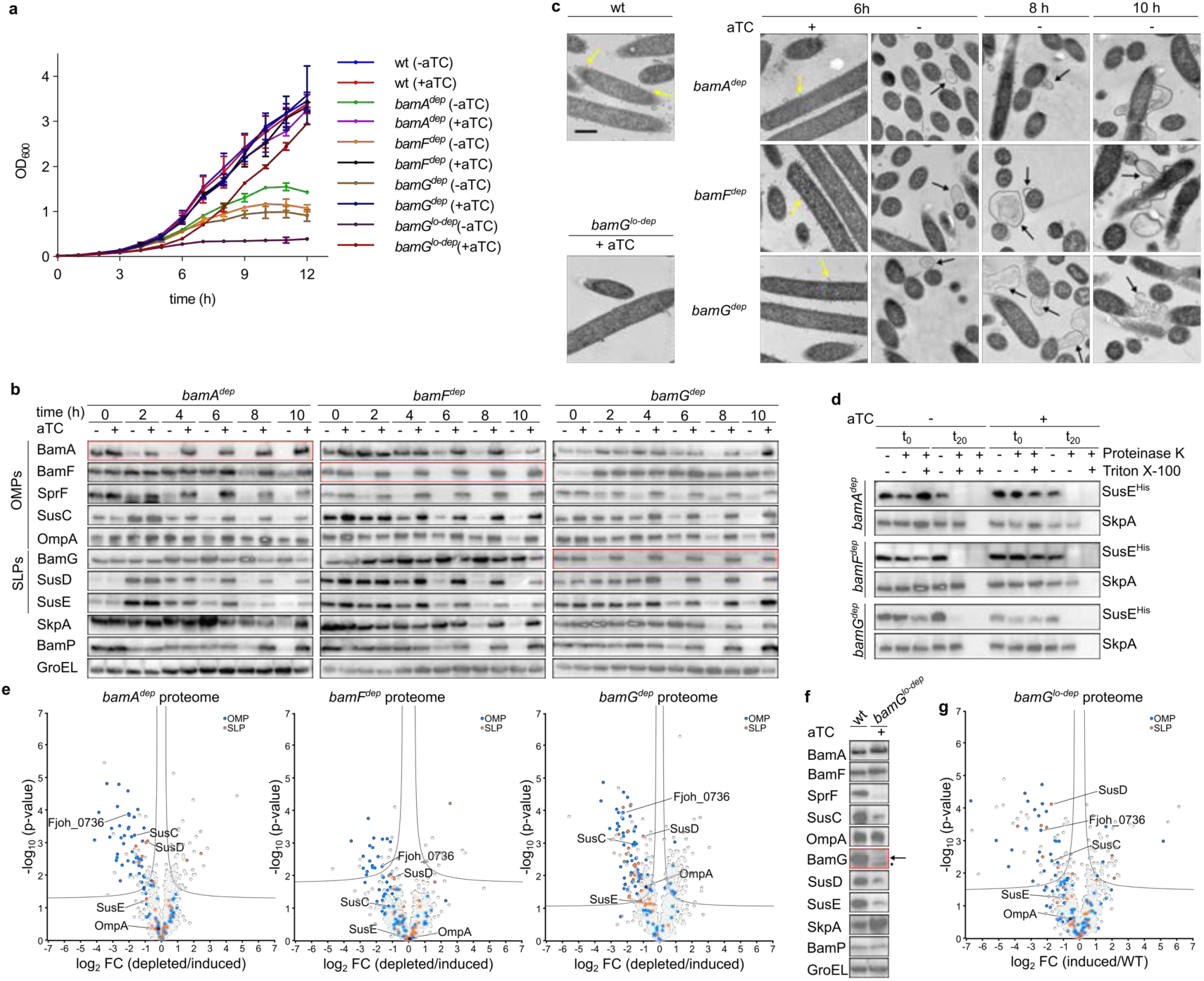
| Depletion analysis of the essential *F. johnsoniae* Bam complex subunits. Strains are the wild type (wt, XLFJ_1078) and corresponding depletion strains for BamA (*bamA^dep^*, XLFJ_1129), BamF (*bamF^dep^*, XLFJ_1115), and BamG with either a strong (*bamG^dep^*, XLFJ_1140) or weak (*bamG^lo-dep^*, XLFJ_1130) inducible promoter. **a-d**, Time course of subunit depletions. The indicated strains were cultured in rich (CYE) medium. The aTC inducer of the target gene was removed (-aTC) at t=0 h where indicated to initiate subunit depletion. Samples in **b-d** were taken at the indicated time points in **a**. Similar data were obtained for three biological repeats. **a**, Growth curves. Shown are the means ± 1 SD. **b**, Whole cell immunoblots. BamA and BamF are detected via epitope tags. SkpA is a periplasmic protein to control for OM integrity. GroEL is a cytoplasmic loading control. The blots for the depleted subunit are boxed in red. **c**, Representative transmission electron microscopy images showing OM defects in the depletion strains. wt and *bamG^lo-dep^* strains were sampled at the 6 h time point in **a**. Yellow arrows highlight budding OM vesicles. Black arrows highlight OM blebbing and rupture. Scale bar, 500 nm. **d**, Depletion of Bam_Fj_ subunits does not change the surface exposure of the SLP SusE. Strains expressing a protease-sensitive His-tagged variant of SusE (SusE^His^, see Extended Data Fig. 7h) were recovered at 6 h after initiation of depletion and treated as indicated with Proteinase K and the detergent Triton X-100 (to permeabilise the OM). Reactions were stopped immediately (t_0_) or after 20 min (t_20_) and analysed by immunoblotting with His tag antibodies. The periplasmic protein SkpA serves as an OM integrity control. Similar results were obtained from 3 biological repeats. **e**, Comparative whole membrane proteome analysis of depleted (-aTC) versus induced (+aTC) for strains harvested at the 6 h time point in **a**. Data points for OMPs and SLPs are coloured as indicated. The data points corresponding to the most highly expressed OM proteins (OmpA, SusCDE, Fjoh_0736) are labelled. A significance threshold is drawn according to t-test parameters for a FDR of 0.1 and a S_0_ of 0.1. Data are averaged over three biological repeats. **f,g** Phenotypic analysis of chronic BamG depletion in an induced strain (*bamG^lo-dep^* + aTC) in which a weak promoter results in the incomplete restoration of wild type BamG levels. Similar results were obtained for 3 biological repeats. **f**, Whole cell immunoblots. Arrow, BamG; *, non-specific band. **g**, As in (**e**) but a comparison of chronic BamG depletion (*bamG^lo-dep^* strain +aTC) relative to the wt strain.

The effects of the BAM subunit depletions on the cellular levels of the remaining BAM_Fj_ components and of representative OMPs and SLPs was assessed by immunoblotting (Fig. 4b). The analysed proteins include the two most abundant *F. johnsoniae* OM components^32,37,38^ namely OmpA (Fjoh_0697), which is an 8-strand OMP that anchors the OM to the cell wall, and a SUS complex of unknown function that we show here to be composed of a 22-strand SusC OMP (Fjoh_0403) together with its SusD SLP partner (Fjoh_0404) and a structurally unrelated SLP SusE (Fjoh_0405)(Extended Data Fig. 7f). The levels of SprF, a 14-strand OMP involved in gliding motility^39^, were also assessed. The effects of depleting all three BAM_Fj_ subunits were broadly similar. OMP levels fell after depletion of the target subunit, although at differing rates. OmpA is notably slow to deplete. It is possible that in addition to BAM_Fj_ other members of the Omp85 family present in the *F. johnsoniae* OM may also be able to insert this simple OMP as has recently been demonstrated for the *E. coli* TAM complex^40^. The levels of the SLPs also decreased (SusD and SusE) with the exception that BamG levels actually increased.

Because *F. johnsoniae* is able to release OM vesicles (Fig. 2c and ^35^) we investigated whether the reduced OM protein levels in the depletion strains were a consequence of OM loss through vesicle shedding. However, no increase in OM protein was detected in the vesicle fraction of the culture supernatant (Extended Data Fig. 7g). Thus, as in *E. coli*^41^, the OM is not being lost through vesicle production when BAM is depleted. The observed reduction in OMP levels therefore reflects defects in their biogenesis.

Analysis of the surface exposure of the SLP SusE provides no evidence that SLPs are accumulating inside the depletion strains and thus no evidence that their export is blocked (Fig. 4d, Extended Data Fig. 7h).

We extended our analysis of the effects of the Bam subunit depletions to the whole OM proteome (Fig. 4e, Extended Data Fig. 7i, Supplementary Data 1). We analysed membranes collected 6 hours after removal of the inducer at which point depletion of the target subunit is complete but the other BAM_Fj_ subunits are still present and the OM is still intact (Fig. 4a-c). The overall pattern of OM proteome changes in all three depletions is similar with marked decreases in the levels of many OMPs (Fig. 4e and Extended Data Fig. 8a). A number of SLPs also decrease in abundance. Thus, removal of the essential BAM_Fj_ subunits has the general effect of reducing the levels of OM proteins.

As an alternative to fully depleting the BAM_Fj_ subunits, we also investigated the effects of chronically reducing the steady state concentration of BamG to a level at which there is a marked effect on cell growth (Fig. 4a,f). Cells of this strain had less severe defects in OM morphology than after full BamG depletion although the budding of OM vesicles seen in the parental strain was almost fully suppressed (Fig. 4c). The differences in the steady state OM proteome in this strain relative to that in wild type cells followed the same trends as the proteome changes seen in the total depletion experiments in showing a general reduction in OMPs and SLPs (Fig. 4f,g and Extended Data Figs. 7i and 8a).

In summary, the loss of either BamF or BamG results in changes in the OM proteome and cellular morphology that closely match those associated with the total loss of BAM function that occurs when BamA is removed. Thus, BamF and BamG are both essential for the core BAM_Fj_ function of OMP insertion.

## Isolation and characterization of a *bamG* suppressor mutant

The requirement for BamF and BamG in BAM_Fj_ function could reflect a direct involvement of these subunits in the general OMP biogenesis function of the BAM complex. However, the same phenotype could also arise indirectly if BamF and BamG have a specialised role in the maturation of a subset of BAM_Fj_ clients such that in their absence these clients accumulate on BamA and interfere with its ability to carry out general OMP biogenesis. To test this second possibility, we asked whether substrate proteins were stably trapped on BamA complexes isolated from strains depleted for BamF or BamG or, as a control, deleted for BamM. In each case a BamADP or BamADFP complex was recovered (Extended Data Fig. 9a,b) indicating that linking BamF and BamM together via BamG stabilises their interactions with BamA. However, no additional proteins that could correspond to trapped substrate molecules were co-purified with any of the preparations.

If the hypothesis that BamF and BamG have client-specific roles in BAM_Fj_ function was correct, then we envisaged that it might be possible to identify suppressor mutations that relieve the secondary effects of BamF/G removal on general BAM function. We were able to select a spontaneous mutant of the BamG depletion strain that allowed growth in the absence of the inducer aTC. Genome sequencing identified a Q801K substitution in BamA as most likely to be responsible for the suppressor phenotype. Re-creation of the BamA Q801K substitution in a clean background permitted deletion of both *bamG* and its orthologue *bamG2* confirming that this single amino acid substitution was responsible for the *bamG* suppressor phenotype. The resultant *bamA^Q801K^*Δ*bamG* Δ*bamG2* strain (hereafter *bamG^sup^*) grew as rapidly as the wild type strain on rich medium (Fig. 5a) even though BamG is absent (Fig. 5b). Thus, although *bamG* behaves as an essential component of BAM_Fj_ in the native context, it is dispensable in an experimentally-modified genetic background. This has parallels to the way that *E. coli* BamD can be deleted in a *bamA* suppressor background^42^. Importantly, the *bamA^Q801K^* mutation did not allow deletion of *bamF,* indicating that BamG and BamF have non-identical functions (Extended Data Fig. 4b).

**Fig. 5.**
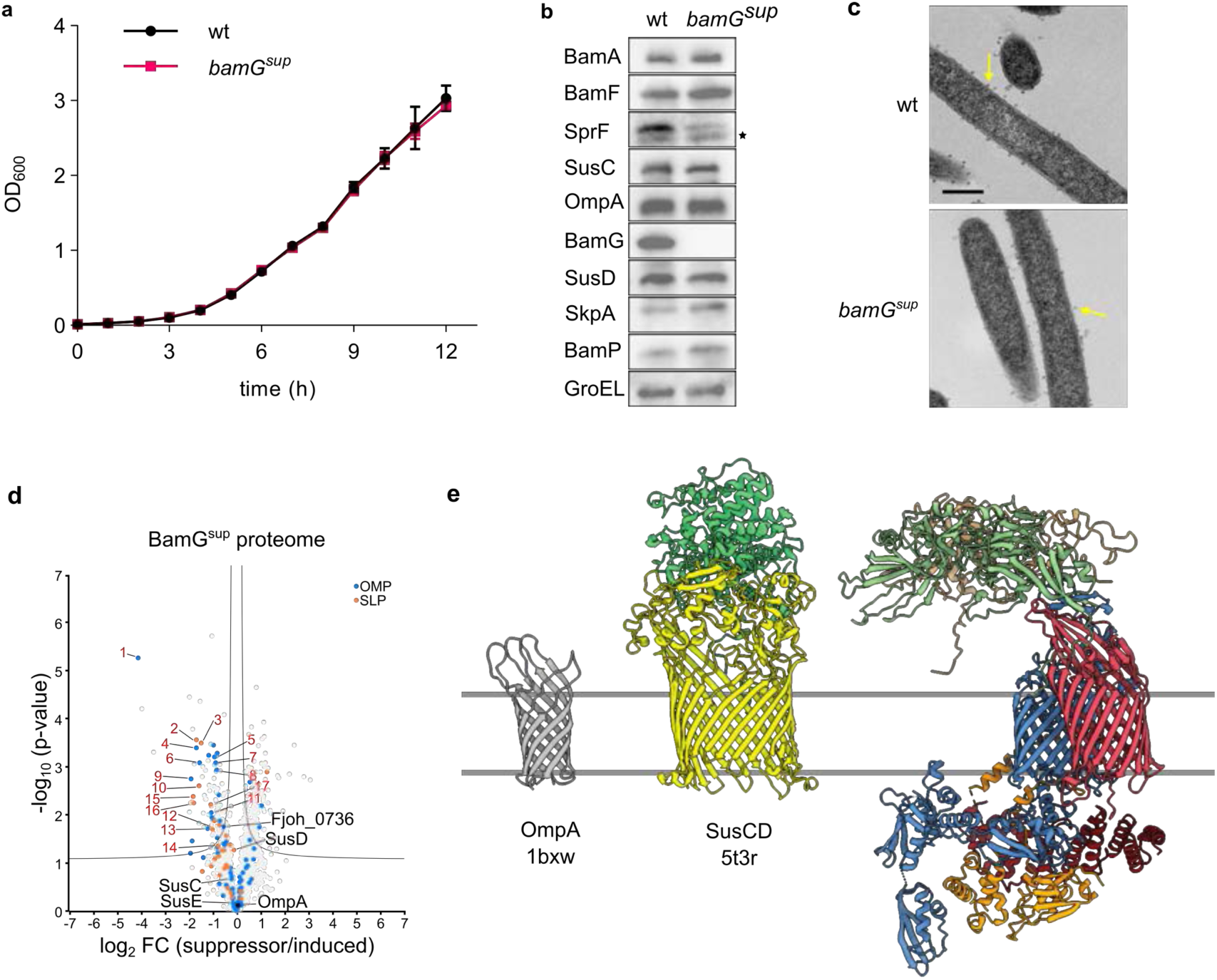
| Characterisation of a *bamG* suppressor mutant. Comparative characterization of the recreated *bamG^sup^* mutant (*bamA^Q801K^* Δ*bamG* Δ*bamG2*, strain XLFJ_1198) and wild type (wt, strain XLFJ_1078) strains. **a,b,d,e,f** Similar data were obtained for three biological repeats. Cells were analysed (**b,e**) and membranes prepared (**c,d,f**) at the 6 h time point in **a**. **a**, Growth on rich (CYE) medium in the absence of aTc. Error bars show the mean ± 1 SD. **b**, Whole cell immunoblots. SkpA is a periplasmic protein to control for OM integrity. GroEL is a cytoplasmic protein as loading ocy. BamA and BamF are detected via epitope tags. *, non-specific bands. **c**, Representative transmission electron microscopy images of the wt and *bamG^sup^* mutant. Yellow arrows highlight budding OM vesicles. Scale bar, 500 nm. **d**, Comparative whole membrane proteome analysis of the *bamG^sup^* strain relative to a BamG-induced strain (*bamG^-dep^* +aTC). Data points for OMPs and SLPs are coloured as indicated. The data points corresponding to the most highly expressed OM proteins (OmpA, SusCDE, Fjoh_0736) are labelled in pink. Proteins that show poor recovery in the *bamG^sup^*strain in a post hoc ANOVA analysis with BamG-induced and depleted strains are numbered as in Extended Data Fig. 8b. A significance threshold is drawn according to t-test parameters for a FDR of 0.1 and a S_0_ of 0.1. Data are averaged over three biological repeats. **e**, Size comparison between BAM_Fj_ and a typical SusCD complex and OmpA. SusCD and OmpA are illustrated using homologous proteins of known structure from other organisms (labelled with the PDB accession numbers).

The *bamG^sup^* strain had normal cellular morphology (Fig. 5c) and no defect in OM integrity, SLP export, or gliding motility (Extended Data Fig. 10a-c). The levels of SusC, SusD, and SusE were restored to wild type levels (Fig. 5b) and the cell was able to assemble these proteins into SusCDE complexes (Extended Data Fig. 10d). Thus, the most abundant *F. johnsoniae* SUS system does not rely on BamG for its biogenesis.

Analysis of the OM proteome of the suppressor strain showed strong restoration of the levels of many OMPs and SLPs relative to BamG-depleted conditions (Fig. 5d and Extended Data Fig. 8b). However, the levels of other OM proteins were poorly recovered suggesting that these proteins were particularly sensitive to the loss of BamG. These sensitive proteins were almost all SusCD pairs and their SLP partners (Extended Data Fig. 8b). Thus, BamG may be particularly important in the biogenesis of a subset of SUS systems.

## Discussion

The BAM complex from the Bacteroidota *F. johnsoniae* is extensively remodelled and elaborated relative to the canonical BAM complex of the γ-Proteobacterium *E. coli* (Fig. 1c,d). *F. johnsoniae* BAM retains only the BamAD core from the *E. coli* complex, dispensing with the other periplasmic lipoprotein subunits, but adding novel subunits located in the OM (BamF), in the extracellular environment (BamG and BamM), and inserted into the BamA barrel (BamP). Of these additional subunits only BamF and BamG are universally conserved across the Bacteroidota (Fig. 3a), suggesting that these proteins are central to the operation of the Bacteroidota BAM system. This inference is supported by our observation that these two subunits, but not BamM and BamP, are essential for cell survival in an otherwise wild type background. Thus, the essential core of BAM has been expanded from BamAD in the canonical BAM_Ec_ complex to BamADFG in Bacteroidota BAM.

Depletion of the central BamA subunit reveals the cellular effects of loss of BAM_Fj_ function (Fig. 4a-c). As expected, the major impact is on OMP biogenesis. However, levels of SLPs also decrease to a lesser extent. Whilst this would be consistent with a role for BAM_Fj_ in SLP biogenesis, assigning causality is complex because any OM protein involved in this process will indirectly depend on BAM_Fj_ for their own biogenesis. Furthermore, our biochemical analysis provides no evidence that loss of BAM_Fj_ leads to the accumulation of non-exported SLPs inside the cell (Fig. 4d).

During our BAM_Fj_ subunit depletion experiments a considerable reduction in the level of the target subunit is required before there is an effect on growth rate or OMP biogenesis (Fig. 4a,b,f). Similarly, even under inducing conditions the levels of BamG in the BamG depletion strain are only small fraction of those in the wild type bacterium (Extended Data Fig. 7d) yet growth in rich medium and OM morphology are unaffected (Fig. 4a,c and Extended Data Fig. 7e). These observations, indicate that the BAM_Fj_ system has considerable excess capacity even in rapidly growing cells. This may explain why the BamM and BamP subunits, if they promote BAM catalytic efficiency rather than being mechanistically essential, can be removed without obvious phenotypic effect when cultured in the laboratory under either rapid (rich medium) or slow (single sugar carbon source in minimal medium) growth conditions (Extended Data Fig. 4c,e).

The novel BamF, BamG, and BamM subunits of BAM_Fj_ form a canopy structure at the extracellular side of the complex and anchor this to BamA. This canopy provides a protected extracellular cavity above the position in the membrane where client OMPs assemble on BamA and suggests that it functions to form an extracellular folding vestibule. Consistent with this hypothesis, BamM has a PPI-like domain that could assist in folding. A possible precedent for folding assistance at the *trans* side of an BAM-like machine comes from the mitochondrial SAM complex which contains subunits that contact client OMPs from the cytoplasmic (external) side of the membrane^43^. The BAM canopy might protect folding intermediates on the BAM complex from proteolysis through sterically blocking the access of proteases present in the extracellular environment. Similarly, the presence of the canopy should sterically exclude LPS molecules (which have large head groups and form rigid arrays in the OM^1,44^) from the region of the membrane next to the lateral seam. This would provide a patch of phospholipid bilayer in the OM for client OMPs to fold into.

The novel BamF, BamG, and BamM subunits of BAM_Fj_ are likely to expand the range of OMPs inserted relative to the canonical BAM_Ec_ complex and, therefore, act on specific structural classes of proteins that are only found in the Bacteroidota. In addition, their cell surface location implies that the novel subunits act on the extracellular portions of BAM substrates. Given these expectations it most likely that these new components are involved in one or more of the following processes: biogenesis of OMPs with large extracellular regions; assisting BamA to transport and fold SLPs (but see comments above); allowing the assembly of the abundant SusCD family complexes that characterise the Bacteroidota OM. In this context it may be significant that the only phylum within the FCB grouping that lacks BamF and BamG proteins (the Chlorobiota) also lacks both SusCD systems and the FCB-specific T9SS translocon channel SprA, which has more than 150kDa of polypeptide on the extracellular side of the membrane^20^ (Fig. 3a). Thus there is a correlation between having a BAM_Fj_-like BAM complex and being able to build the SusCD systems and T9SS translocon that characterize the FCB superphylum. Our observation that a subset of SusCD proteins are only minimally recovered by a *bamG* suppressor mutation (Fig. 5d) supports the idea that at least BamG is involved in the assembly of some SusCD systems.

In Fig. 5e we compare the proportions of BAM_Fj_ with those of the SusCD unit that it may assemble as well as the more classical *E. coli* OMP substrate OmpA that does not have an extensive extracellular domain. What is immediately obvious is that whilst SusC can be accommodated under the BAM_Fj_ canopy, the full SusCD complex cannot do so without the canopy being raised. However, displacement of the canopy appears unlikely due to the tethering of the canopy to BamA and BamF at one end, and to the membrane by the lipid anchor of BamM at the other. If lifting of the BamM anchor end is possible, then this would require either distortion of the membrane bilayer around the lipid anchor and/or unfolding of the first part of BamM. These considerations suggest that SusC is likely to fold on BAM_Fj_ and be at least partially released before forming a complex with its SusD partner.

We were able to select a suppressor mutation in *bamA* that compensates for the loss of the BamG subunit. This single amino acid substitution in BamA is sufficient to restore OMP biogenesis and OM morphology (Fig. 5b,c) showing that general OMP insertion in *F. johnsoniae* does not physically require the presence of BamG. It is unlikely that the suppressing amino acid substitution in BamA functions by replicating the role of BamG since it is difficult to see how alterations in BamA could create a similar structural environment to the BamG-containing extracellular canopy. Instead, it is most plausible that the suppressor substitution acts by compensating for the toxic consequences of loss of BamG function. Since removal of BamG closely phenocopies the loss of BamA (Fig. 4b,c,e-g), the most probable suppression scenario is that loss of BamG blocks BamA function through the accumulation of stalled BamG-requiring substrates and that this blockage is relieved by a structural change in BamA that corrects the problem, for example by accelerating substrate release. The BamA residue that is substituted in the *bamG* suppressor, Gln801, is located in barrel loop 11-12 that lies over the extracellular end of the BamA pore (Fig. 2a). Gln801 is hydrogen-bonded through its side cain oxygen atom to the main chain amine of Gly591 in adjacent loop 9-10 (Fig. 2a) so it is likely that its substitution disrupts the packing of the BamA extracellular cap. We speculate that this may marginally destabilise BAM-substrate interactions allowing the release of malfolded substrates.

Although analysis of the *bamG* suppressor allowed us to identify certain SusCD proteins that are heavily dependent on BamG for their biogenesis, many other SusCD systems, including the most abundant SusCDE complex, were well-restored in the same background (Fig. 5b,d Extended Data Fig. 9d). We interpret this as meaning that most BamG clients are able to fold without BamG during the vast majority of BAM_Fj_ turnovers and that BamG is only required to correct a small proportion of insertion events that go wrong. In this model BamG has a quality control role that prevents infrequent errors in folding blocking the BAM_Fj_ complex. Alternatively BamG may play a more critical role in the biogenesis of these proteins under specific conditions such as stress or under conditions that are not readily replicated in the laboratory.

It is not possible to delete *bamF* in the *bamG*-suppressing background indicating that BamF has non-identical roles(s) to BamG. Since BamF anchors BamG to BamA, loss of BamF most likely affects both BamG function (which should be correctable by the suppressor mutation) and an additional function affecting core BAM OMP insertion activity (which the suppressor cannot correct). Thus, BamF may be an obligate mechanistic partner of BamA in the Bacteroidota.

Our work shows that the Bacteroidota contain a heavily modified BAM complex that presents a new paradigm for outer membrane protein biogenesis in this very important group of organisms.

## Acknowledgments

We thank Mark McBride and Satoshi Shibata for providing antibodies used in this study and Frédéric Lauber for producing the SkpA antiserum. We acknowledge the use of the University of Oxford Department of Biochemistry Advanced Proteomics Facility, the Oxford Micron Advanced Imaging Facility, and the Sir William Dunn School of Pathology Electron Microscopy Facility. We thank Vaishnavi Ravikumar for collecting the proteomics data, and Marjorie Fournier, Faraz Mardakheh, Shabaz Mohammed, and James Holder for advice on interpreting this data, as well as Raman Dhaliwal and Charlotte Melia for carrying out the TEM analysis. This work was supported by European Research Council Advanced Award 833713 (B.C.B). This research was supported in part by the Intramural Research Program of the NIH.

## Author Contributions

XL and LOT carried out all genetic and biochemical experiments. JCD and SML collected electron microscopy data and determined all structures. XL, BCB, and SML conceived the project. BCB and SML supervised the project and secured funding. All authors interpreted data and wrote the manuscript.

## Competing interests

The authors declare no competing interests.

## Materials and Methods

### Bacterial strains and growth conditions

All strains and plasmids used in this work are listed in Tables S3 and S4. *F. johnsoniae* was routinely cultured aerobically in Casitone Yeast Extract (CYE) medium ^45^ at 30 °C with shaking. For some physiological studies the cells were cultured in PY2 medium^46^ as indicated below. For experiments testing growth on complex sugars cells were cultured in a 96-well plate in a CLARIOstarPlus plate reader using the modified minimal A medium described in^33^ and containing 0.25 % (w/v) of either Carob galactomannan (Megazyme, CAS Number: 11078-30-1) or tamarind xyloglucan (Megazyme, CAS Number: 37294-28-3) as the sole carbon source. *E. coli* strains were routinely grown aerobically in LB medium at 37 °C with shaking, or on LB agar plates. Where required, 100 µg ml^-1^ erythromycin was used in growth medium for *F. Johnsoniae*. 100 µg ml^-1^ ampicillin or 50 µg ml^-1^ kanamycin were used in growth medium for *E. coli*. A final concentration of 0.2 µg ml^-1^ and 2 µg ml^-1^. aTC (CAY10009542-50 mg, Cambridge Bioscience Ltd) was used as a final concentration of 0.2 µg ml^-1^ (liquid culture) and 2 µg ml (agar plates).

### Genetic constructs

Plasmids were constructed by Gibson cloning^47^ using the primers and target DNA in Table S3. Suicide and expression plasmids were introduced into the appropriate *F. johnsoniae* background strain by triparental mating as previously described^46^. Chromosomal modifications were introduced using the suicide vector pYT313 harboring the counter-selectable *sacB* gene as previously described^48^. All plasmid constructs and chromosomal modifications were confirmed by sequencing.

### Construction of a tightly regulated gene expression system for *F. johnsoniae*

The aTC-inducible systems for the depletion of essential Bam_Fj_ components (Extended Data Fig. 7a) were based on the native *F. johnsoniae ompA* and *fjoh_0824* promoters and contain the 100bp upstream of *ompA* or *fjoh_0824*. Guided by the observations of Lim and co-workers^49^ a *tetO2* site (TetR binding site) was inserted upstream of the conserved –33 motif in these promoters and another *tetO2* site downstream of the conserved –7 motif generating the synthetic promoters *P_ompAinduc_* and *P_fjoh_0824induc_* (Extended Data Fig. 7b). The constructs also contain *tetR* under the control of an additional copy of the constitutive *F. johnsoniae ompA* promoter. The final inducible systems containing the gene to be induced were integrated into the chromosome at an assumed phenotypically neutral site^32,39^ by replacing *fjoh_4538* to *fjoh_4540*.

The designed inducible systems were validated using strains in which a NanoLuc reporter gene^50^ was placed under the control of the chromosomally-integrated aTC-inducible systems (Extended Data Fig. 7c). Overnight cultures of these strains were diluted 1:100 into fresh CYE medium in the absence or presence of 0.2 µg ml^-1^ aTC and cultured for 6 h to mid-exponential phase (OD_600_ ∼0.6). Cells were collected and resuspended in PY2 medium to OD_600_=0.6. A volume of 50 µl of cell resuspension was mixed with 50 µl of reaction solution (48 µl PY2 medium supplied with 2 µl of furimazine (Promega)) in a 96-well plate and the luminescence signal measured in a CLARIOstar^Plus^ plate reader.

Strains to enable the depletion of the essential BAM_Fj_ subunits were constructed by introducing a copy of the target gene under the control of the designed inducible system into the chromosome at the phenotypically neutral site. The native copy of the target gene was then be deleted in the presence of aTC to allow expression of the introduced copy of the gene.

### Purification of BAM_Fj_ and SusCDE complexes

To purify complexes containing Twin-Strep tagged BamA, the relevant strain was cultured for 22 h in CYE medium using 1 l culture volume in 2.5 l flasks. A total culture volume of 12 l was used for sample preparations for structure determination, and 4 l of culture was used for analytical purifications of BAM_Fj_ variants. Cells were harvested by centrifugation at 12,000*g* for 30 min and stored at −20 °C until further use. All purification steps were carried out at 4 °C. Cell pellets were resuspended in buffer W (100 mM Tris-HCl pH 8.0, 150 mM NaCl, 1 mM EDTA) containing 30 μg/ml DNase I, 400 μg/ml lysozyme and 1 mM phenylmethylsulfonyl fluoride (PMSF) at a ratio of 5 ml of buffer to 1 g of cell pellet. Cells were incubated on ice for 30 min with constant stirring before being lysed by two passages through a TS series 1.1 kW cell disruptor (Constant System Ltd) at 30,000 PSI. Unbroken cells were removed by centrifugation at 20,000*g* for 20 min. The supernatant was recovered and total membranes were collected by centrifugation at 230,000*g* for 75 min. Membranes were resuspended in buffer W to a protein concentration of 6.5 mg/ml and solubilized by incubation with 1% (w/v) lauryl maltose neopentyl glycol (LMNG, Anatrace) for 2 h. Insoluble material was removed by centrifugation at 230,000*g* for 75 min. Endogenous biotin-containing proteins were masked by addition of 1 ml BioLock solution (IBA Lifesciences) per 100 ml of supernatant and incubation for 20 min with constant stirring. The solution was then circulated through a Strep-TactinXT 4Flow High Capacity column (IBA Lifesciences) overnight. The column was washed with 10 column volumes (CV) of buffer W containing 0.01% LMNG (buffer WD) and bound proteins were eluted with 6 CV Strep-TactinXT BXT buffer (IBA Lifesciences) containing 0.01% LMNG. The eluate was concentrated to 500 μl using a 100-kDa molecular weight cutoff (MWCO) Amicon ultra-15 centrifugal filter unit (Merck) and then injected onto a Superose 6 Increase 10/300 GL column (Cytiva) previously equilibrated in buffer WD. Peak fractions were collected and concentrated using a 100-kDa MWCO Vivaspin 500 column (Sartorius).

Purification of SusCDE complexes with a N-terminal Twin-Strep tag on SusC was carried out by the same protocol.

### Peptide mass fingerprinting

Samples were excised from Coomassie-stained gels. For whole sample proteomic analysis, SDS-PAGE was carried out only until the sample had fully entered the gel and the protein smear at the top of the gel was excised. Samples were subject to in-gel trypsin digestion and electrospray mass spectrometry at the Advanced Proteomics Facility (University of Oxford, UK).

### Immunoblotting

Immunoblotting was carried out as previously described^24^. Antibodies against BAM subunits, Sus proteins, and SkpA were raised in rabbits against His-tagged recombinant proteins produced using the plasmids listed in Supplementary Table 2. Antiserum against OmpA^37^ was provided by Satoshi Shibata (Tottori University) and antiserum against SprF^39^ by Mark McBride (University of Wisconsin-Milwaukee). The following commercial antisera were used: anti-StrepTag (34850 Qiagen), anti-GroEL (G6532 Merck), anti-Alfa Tag (N1582 Synaptic Systems GmbH), anti-His tag (H1029-100UL Merck Life Science UK Limited), anti-mouse IgG peroxidase conjugate (A4416 Merck), and anti-rabbit IgG peroxidase conjugate (31462 Pierce).

### BAM subunit depletion experiments

The desired depletion strain was grown overnight in CYE medium supplemented with 0.2 µg ml^-1^ aTC. Cells from 1 ml of the overnight culture were collected, washed once in 1ml CYE, and resuspended in 1 ml of CYE medium. Cells from this sample were then used to inoculate 15 ml of CYE medium, either with or without 0.2 µg ml^-1^ aTC, to OD_600_=0.02. The cells were then cultured aerobically at 30 °C and cell samples collected into SDS sample buffer every 2 h for subsequent analysis by immunoblotting. Samples for imaging or membrane preparation were collected and analysed as detailed below.

To purify BamA complexes after depleting the essential BamF or BamG subunits, a 200 ml overnight culture of the appropriate strain grown in the presence of 0.2 µg ml^-1^ aTC was collected and resuspended in the same volume of fresh CYE medium without aTC. This sample was used to inoculation 8 l of CYE without aTC to OD_600_=0.1 which was then cultured aerobically at 30 °C for 6 h. Cells were collected and BamA complexes processed for purification as described above.

### Microscopic analysis of cells during BAM subunit depletions

Live cells were imaged directly in growth medium by spotting samples taken from depletion cultures onto a 1% agarose pad prepared in PY2 medium. Phase contrast images were acquired on an inverted fluorescence microscope (Ti-E, Nikon) equipped with a perfect focus system, a 100× NA 1.4 oil immersion objective, a motorized stage, and a sCMOS camera (Orca Flash 4, Hamamatsu).

For transmission electron microscopy, cells were collected at the required time points during depletion by centrifugation at 8,000 g for 5 min. After carefully removing the supernatant, cell pellets were gently resuspended in 1 ml of fixative solution (2.5% glutaraldehyde, 4% formaldehyde in 0.1 M PIPES buffer, pH 7.4) and incubated at room temperature for 1 h. Following fixation cells were washed with TEM buffer (100mM PIPES NaOH pH 7.2), treated with TEM buffer containing 50 mM glycine, washed again in TEM buffer, and then subjected to secondary fixation with TEM buffer containing 1% (w/v) osmium tetroxide and 1.5% (w/v) potassium ferrocyanide. Samples were then washed extensively with Milli-Q water, stained with aqueous 0.5% (w/v) uranyl acetate overnight, then washed again with Milli-Q water. The samples were dehydrated through an ethanol series and infiltrated with and embedded in TAAB low viscosity epoxy resin ahead of polymerisation at 60 °C for 24 h. Sections of 90 nm were cut from the resin blocks using a Leica UC7 Ultramicrotome and collected onto 3 mm copper grids. The sections were then post-stained with lead citrate and imaged using a JEOL Flash 120kV TEM equipped with a Gatan Rio camera.

### Whole membrane proteomics

15 ml of cells at the 6 h time point of the standard depletion experiment were collected by centrifugation at 8,000 g for 5 min at 4 °C. The cells were resuspended in 1 ml of buffer W and lysed on ice using a probe sonicator (Sonics Vibra Cell, probe 630-0422) at 40% power by 12 repeats of a 10 s on/10 s off pulse cycle. After lysis, the samples were centrifuged at 20,000 g for 20 min at 4 °C to remove cell debris. The supernatant was then centrifuged at 135,000 g for 45 min at 4 °C to pellet the membranes. The membranes were resuspended in buffer W and the protein contents of the samples normalised by A_280nm_. The samples were run together on SDS-PAGE gels and stained with Coomassie Blue (Extended Data Fig. 7j) to confirm that normalisation had been correctly implemented.

Membrane fractions were resuspended in lysis buffer containing 1% SDS, 0.1 M ammonium bicarbonate pH 8.0. Samples were sonicated for 5 x 15s in a water bath with 15 s incubations on ice between each pulse cycle. The samples were clarified by centrifugation at 17,500 g for 30 min and 50 µg of total protein lysate was taken for analysis. Samples were reduced for 30 min using 10 mM tris(2-carboxyethyl)phosphine (TCEP) followed by alkylation for 30 min in the dark using 2-chloroacetamide. SpeedBeads Magnetic Carboxylate Modified Particles (GE Healthcare) were mixed with the sample in a 10 volumes beads: 1 volume sample ratio and the samples shaken for 10 min at 1,000 rpm. The beads were then washed twice (8x volumes) with 70 % ethanol followed by 100 % acetonitrile. 100 mM ammonium bicarbonate was added to the washed beads and pre-digestion with endoprotease LysC (Wako; 1:100) was carried out at 37 °C for 2 h. This was followed by 16 h digestion with trypsin (Promega, 1:40) at 37°C. The supernatant was collected and any remaining bound peptides were eluted from the beads using 2% dimethyl sulfoxide (DMSO). Digested peptides were loaded onto C18 stage tips, pre-activated with 100 % acetonitrile and 0.1 % formic acid and centrifuged at 4000 rpm. The tips were then washed with 0.1% formic acid and eluted in 50 % acetonitrile / 0.1 % formic acid. Eluted peptides were dried in a speed-vac.

Peptide analysis employed a Thermofisher Scientific Ultimate RSLC 3000 nano <u>liquid</u> <u>chromatography</u> system coupled in-line to a Q Exactive mass spectrometer equipped with an Easy-Spray source (Thermofisher Scientific). Peptides were separated using an Easy-Spray RSLC C18 column (75µm i.d., 50 cm length, Thermofisher Scientific) using a 60 min linear 15 % to 35 % solvent B (0.1 % formic acid in acetonitrile) gradient at a <u>flow rate</u> 200 nL/min. The raw data were acquired on the mass spectrometer in a data-dependent acquisition (DDA) mode. Full-scan MS spectra were acquired in the Orbitrap (Scan range 350-1500 *m*/*z*, resolution 70,000, AGC target 3e6, maximum injection time 50 ms). The 10 most intense peaks were selected for higher-energy collision dissociation (HCD) fragmentation at 30 % of normalized collision energy. HCD spectra were acquired in the Orbitrap at resolution 17,500, AGC target 5e4, maximum injection time 120 ms with fixed mass at 180 *m*/*z*.

MS data was analyzed using MaxQuant 2.5.1.0 as previously described^51^ to obtain label-free quantification (LFQ) values that were then used for data processing in Perseus 2.1.3.0^52^. LFQ values were log_2_ transformed and categorically grouped by replicates. Rows were filtered based on 2 valid values in each group and then missing values were replaced using a normal distribution with a width of 0.3 and down shift of 1.8 (default values). Then, dataset was normalized by subtracting the medians of each sample. After visually verifying a normal distribution and a linear correlation, sample pairs were subjected to a two-tailed T test using a FDR of 0.1 and a S_0_ of 0.1 to define a threshold of statistical significancy. Proteins were represented in a volcano plot, according to the log_2_ of their enrichment and the –log_10_ of the T-test p-value.

An ANOVA test was carried out for indicated groups of proteins using a Benjamini-Hochberg method with a FDR of 0.05 for truncation. Then, a post hoc Turkey’s Honest Difference Significance (HDS) test for one-way ANOVA using a FDR of 0.05 was carried out. Proteins were then filtered by ANOVA-significance and by category to represent in a heat map their HDS scores, as indicated.

A batch normalization using empirical Bayes method was carried with the ComBat script^53^ for PerseusR package^54^ to make the heat map for all depletions (Fig. ED 9). Then, samples were subjected to the statistical test previously described.

The proteins obtained from the MS experiments were categorised as follows. Proteins with signal peptides or lipoprotein signal peptides were first extracted using SignalP 6.0^55^ to obtain datasets containing only OM plus periplasmic proteins, or lipoproteins, respectively. Proteins were then manually sorted to the categories OMP or SLP. This sorting was carried out using Uniprot entry data that included AlphaFold^28^ models. Lipoproteins were only classified as SLPs if they were either SusD homologues or if they were found at a locus coding SusCD systems.

### Determination of cell surface exposure of SusE

The strain for analysis was transformed with plasmid pXL184 which expresses His-tagged SusE. The cells were then grown overnight in CYE supplemented with erythromycin, and for BAM subunit depletion strains with 0.2 µg ml^-1^ aTC. Cells were collected, resuspended in CYE medium, and then used to inoculate 10 ml of erythromycin-containing CYE medium to OD_600_=0.02, supplementing with 0.2 µg ml^-1^ aTC as required. The cells were cultured for 6 h before being collected by centrifugation and resuspended in phosphate buffered saline (PBS) containing 10 mM MgCl_2_ to a total volume of 80 µl and OD_600_=1. Samples were supplemented as appropriate with 200 μg ml^−1^ proteinase K (ThermoFisher) and 1% (v/v) Triton X-100 (Merck) and incubated for 20 min at room temperature. Reactions were stopped by the addition of 5 mM phenylmethylsulphonyl fluoride (ITW Reagents) followed by incubation at 100 °C for 5 min, addition of SDS-PAGE sample buffer, and further incubation at 100 °C for 5 min before analysis by immunoblotting.

### Isolation of outer membrane vesicle fraction

The isolation of outer membrane vesicles (OMVs) was performed essentially as in ^41^. Briefly, cells were separated from culture supernatant by centrifugation at 8,000 g for 5 min and the pellets reserved as the whole cell fraction. Culture supernatant from the equivalent of 2 ml of culture at OD_600_=1 was filtered through a 0.2 µm filter (MilliporeSigma, Cat. #SLGPR33RB) and concentrated using a 100 kDa molecular weight cut-off Amicon Ultra-4 centrifugal filter (MilliporeSigma, UFC810096) to produce the OMV fraction. Samples were adjusted to equal volume before analysis by immunoblotting.

### Isolation of a spontaneous suppressor of BamG depletion

The BamG depletion strain XLFJ_1140 was grown overnight in CYE medium supplied with aTC. 1 ml of cells was collected by centrifugation at 8,000g for 3 min, washed once with CYE and then diluted to a starting OD_600_=0.2 in 10 ml fresh CYE medium without aTC. After culturing for 6 h, cells were diluted 1:200 into fresh CYE medium without aTC and cultured for a further 2 days before plating on CYE agar to obtain single colonies. Individual clones were cultured in parallel with and without aTC in CYE and the expression of BamG analysed by whole cell immunoblotting. Clones that grew without aTC but still expressed BamG only following aTC induction (showing that they were not constitutively de-repressed for BamG synthesis) were subjected to genome sequencing (Plasmidsaurus). This identified the potential suppressor mutation *bamA(Q801K)* which was introduced into a BAM wild-type background, followed by successive deletions of *bamG* and *bamG2* to produce the *bamG^sup^* strain XLFJ_1198.

### Cryo-EM sample preparation and imaging

4 μl of either fraction A (for the BAM_Fj_ complex,1.3 mg/ml) or fraction B (for the BamAP complex, 1.3 mg/ml) of the BAM_Fj_ preparation (Fig. 1a), or of the BamP-deleted BAM complex (ΔBamP complex, 1.2 mg/ml) was adsorbed onto glow-discharged holey carbon-coated grids (Quantifoil 300 mesh, Au R1.2/1.3) for 10 s. Grids were blotted for 2 s at 10 °C, 100% humidity and frozen in liquid ethane using a Vitrobot Mark IV (Thermo Fisher Scientific).

Movies were collected in counted mode, in Electron Event Representation (EER) format, on a CFEG-equipped Titan Krios G4 (Thermo Fisher Scientific) operating at 300 kV with a Selectris X imaging filter (Thermo Fisher Scientific) and slit width of 10 eV, at 165,000x magnification on a Falcon 4i direct detection camera (Thermo Fisher Scientific), corresponding to a calibrated pixel size of 0.732 Å. Movies were collected at a total dose ranging between 52.0-60.3 e^-^/Å^2^ (Table 1), fractionated to ∼ 1.0 e^-^/Å^2^ per fraction for motion correction.

### Cryo-EM data processing

Patched motion correction, CTF parameter estimation, particle picking, extraction, and initial 2D classification was performed in SIMPLE 3.01^56^. All downstream processing was carried out in cryoSPARC2^57^ or RELION 4.03^58^, using the csparc2star.py script within UCSF pyem4^59^ to convert between formats. Global resolution was estimated from gold-standard Fourier shell correlations (FSCs) using the 0.143 criterion and local resolution estimation was calculated within cryoSPARC.

The cryo-EM processing workflow for the BAM_Fj_ complex is outlined in Exended Data Fig. 1. Briefly, particles were subjected to one round of reference-free 2D classification (k=200) using a 240 Å soft circular mask within cryoSPARC resulting in the selection of 2,153,927 clean particles. A subset of these particles (180,179) was subjected to multi-class *ab initio* reconstruction using a maximum resolution cutoff of 7 Å, generating four volumes. These volumes were lowpass filtered to 20 Å and used as references in a heterogeneous refinement against the full 2D-cleaned particle set. Particles (903,299) from the most populated and structured class were selected and non-uniform refined against their corresponding volume lowpass-filtered to 15 Å, generating a 3.0 Å map. Bayesian polishing in RELION followed by duplicate particle removal generated a 2.5 Å map after non-uniform refinement, which could be further improved to 2.3 Å after local and global CTF refinement (fitting beam tilt and trefoil only). These particles were then subjected to heterogeneous refinement against four compositionally distinct volumes previously generated by RELION 3D classification (k=8, 3.75° sampling) of a particle subset of pre-polished particles. Particles (274,708) belonging to the class with strong BamD and POTRA densities were selected and non-uniform refined against their corresponding volume, generating a 2.4 Å map. Additional alignment-free 3D classification in RELION was performed (k=6) using a soft mask covering BamD and the BamA POTRA domains yielding a class with stronger density. Particles (55,795) from this class were selected and non-uniform refined against a previous volume lowpass filtered to 15 Å, generating a consensus 2.7 Å volume. Local refinements were performed against the consensus volume (lowpass filtered to 7 Å) using soft masks covering the BamD/POTRA domains or extracellular density, yielding 3.2 Å and 2.7 Å volumes, respectively. ChimeraX^60^ was used to generate a composite map from the consensus and individual focused maps.

The cryo-EM processing workflow for the BamAP complex is outlined in Extended Data Fig. 6. Two datasets were collected for this sample. In the first dataset particles were subjected to two rounds of reference-free 2D classification (k=200) using a 200 Å soft circular mask resulting in the selection of 979,474 clean particles. These particles were then subjected to multi-class ab initio reconstruction (k=4) using a maximum resolution cutoff of 8 Å, generating four volumes. Particles (514,326) belonging to the two most prominent volumes were combined and non-uniform refined against one of their corresponding volumes, lowpass-filtered to 15 Å, generating a 3.7 Å volume. The second particle dataset underwent four rounds of 2D classification (k=200, 200 Å soft circular mask) followed by multi-class ab initio reconstruction using a maximum resolution cutoff of 7 Å, generating six volumes. Particles (438,412) from the most populated class were selected and refined against their corresponding volume lowpass-filtered to 15 Å, generating a 3.7 Å volume. Particles from both datasets were independently polished within RELION, combined, and non-uniform refined, fitting per-particle CTF parameters, yielding a 3.5 Å map. Alignment-free 3D classification was subsequently performed within cryoSPARC (k=6), using a soft mask covering the full protein density of the complex. Particles (96,076) from the class demonstrating strong density for the N-terminal domain of BamP were selected and non-uniform refined against their corresponding volume, lowpass filtered to 15 Å, generating a 3.7 Å map.

The cryo-EM processing workflow for the ΔBamP complex is outlined in Extended Data Fig. 5. Briefly, particles were subjected to two rounds of reference-free 2D classification (k=200) using a 180 Å soft circular mask within cryoSPARC resulting in the selection of 1,177,554 clean particles. These particles were then subjected to multi-class *ab initio* reconstruction using a maximum resolution cutoff of 6 Å, generating six volumes. Particles from volume classes containing BamA barrels were independently non-uniform refined against their corresponding volume, lowpass filtered to 15 Å. These particles were subsequently combined and refined against a volume (lowpass filtered to 15 Å) from the most populated class, generating a 3.6 Å consensus volume. Bayesian polishing in RELION followed by non-uniform refinement and fitting of per-particle CTF parameters plus beam tilt and trefoil generated a 3.5 Å map. Map quality was further improved by non-uniform refinement of a cleaner particle set (534,368 particles) generated by an additional round of 2D classification (k=100, 180 Å soft circular mask), despite no increase in nominal resolution. A second β-barrel could be resolved in map density at low contour level (0.08). Attempts to improve map quality for this partner β-barrel, through extensive 3D classification and local refinement schemes, did not improve map quality for this region.

### Model building, structure refinement, and figure preparation

Iterative model building and real-space refinement using secondary structure, rotamer, and Ramachandran restraints was performed in Coot v0.9^61^ and Phenix^62^, respectively. Validation was performed in Molprobity^63^ within Phenix. Cryo-EM data collection, image processing and structure refinement statistics are listed in Supplementary Table 1. Figures were prepared using UCSF ChimeraX v.1.4^60^.

## Data availability

Cryo-EM density maps and atomic coordinates are deposited in the Electron Microscopy DataBank (EMDB) with the following accession numbers: EMD-48835 (BAM_Fj_ composite map), EMD-48832 (BAM_Fj_ consensus map), EMD-48833 (BAM_Fj_ BamGM-focused map), EMD-48834 (BAM_Fj_ BamADP-focused map), EMD-48836 (BamAP complex), EMD-48837 (BamAD complex). Atomic coordinates are deposited in the Protein Data Bank (PDB) with the following accession numbers: 9N2D (BAM_Fj_ complex), 9N2E (BamAP complex), 9N2F (BamAD complex).

Requests for materials should be addressed to BCB.

**Extended Data Fig. 1.**
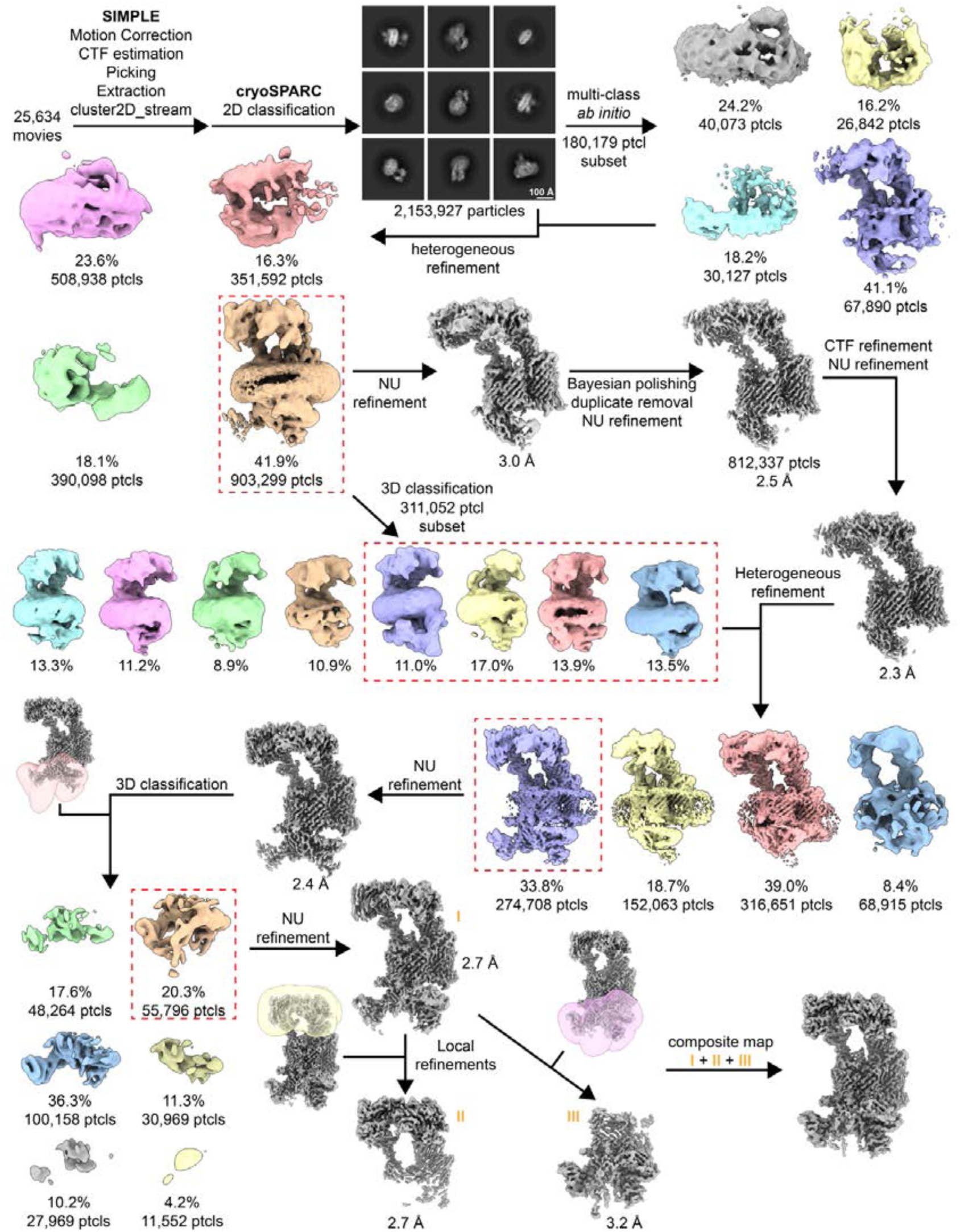
| Workflow for the cryoEM analysis of the *F. johnsoniae* BAM_Fj_ complex. TwinStrep-tagged BamA complexes were purified by Streptactin affinity chromatography and size exclusion chromatography and the major (highest molecular size) peak was analyzed. See Fig. 1a for corresponding SDS-PAGE analysis of this material. Image processing workflow for the BamA complexes.

**Extended Data Fig. 2.**
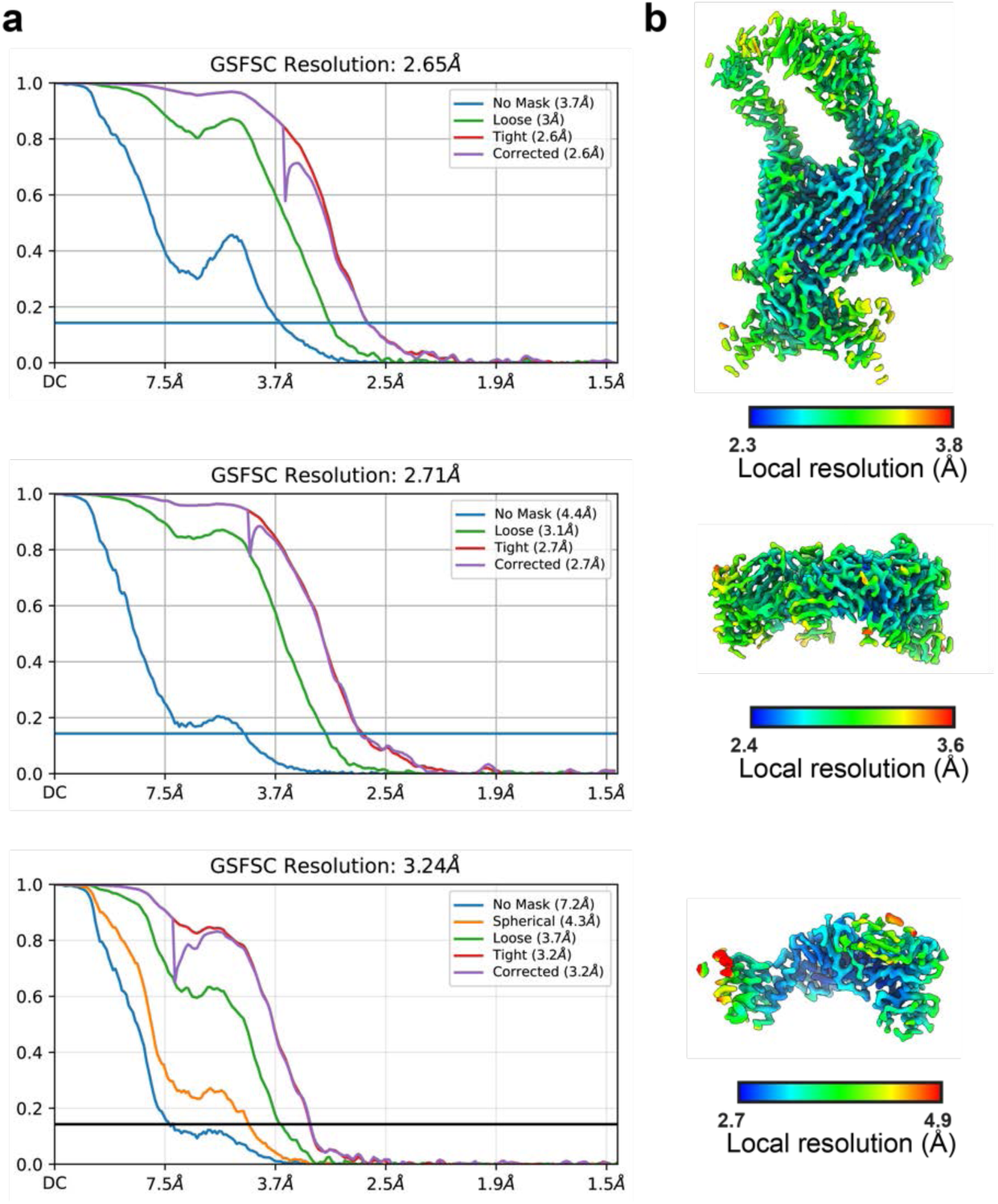
| Map quality metrics for the BAM_Fj_ complex. Gold-standard Fourier Shell Correlation (FSC) curves used for global resolution estimation (**a**) and local resolution estimate (**b**) of consensus (top), extracellular (middle), or periplasmic (bottom) volumes from the BAM_Fj_ complex.

**Extended Data Fig. 3.**
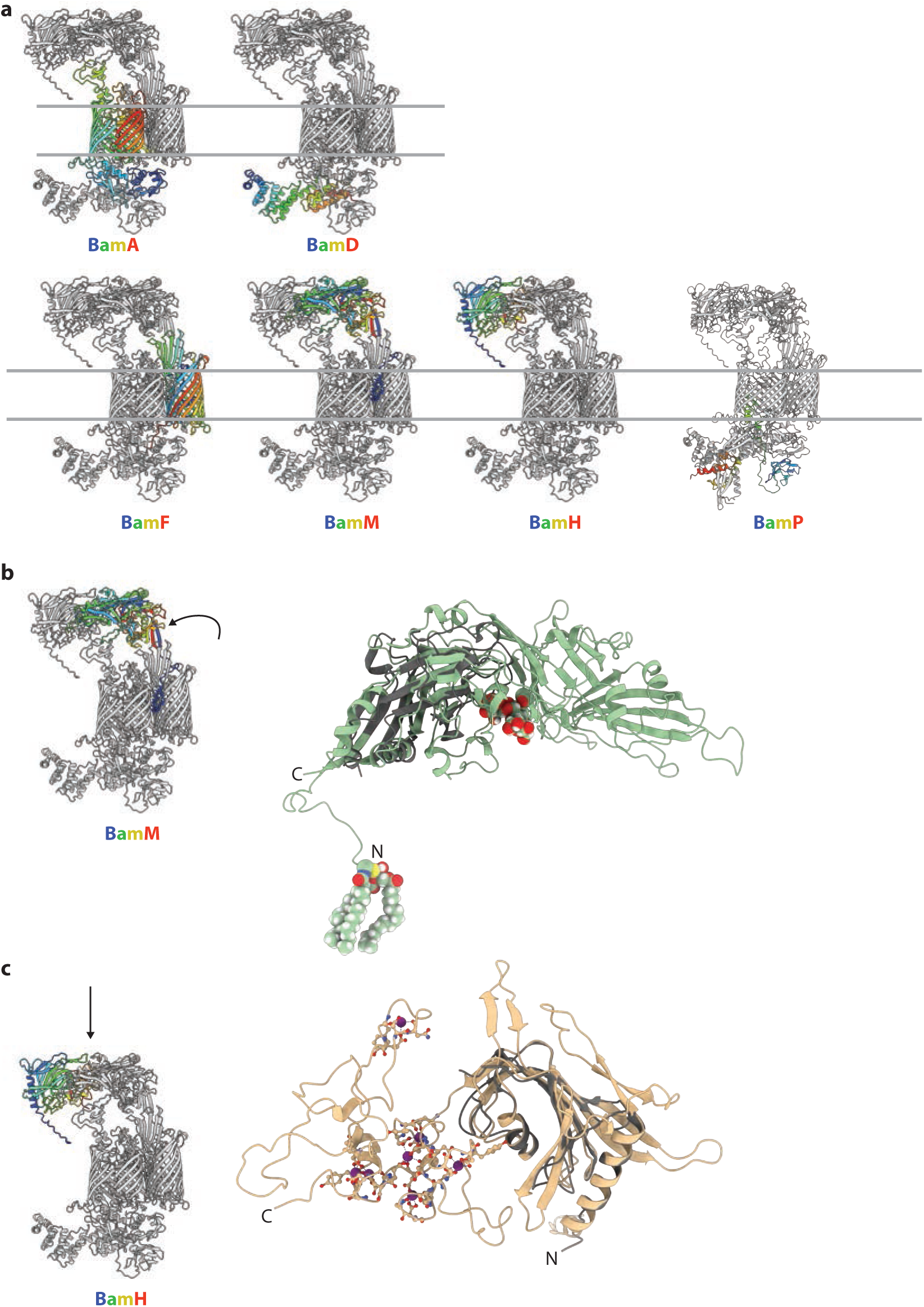
| Further structural analysis of the BAM_Fj_ complex. **a**, Chain ordering. The indicated subunit in each panel is rainbow-coloured from the N-(blue) to C-terminus (red). **b**, BamG (green) in cartoon representation viewed from the direction indicated on the Left Hand model and overlaid with the closest structural homologue, the chondroitin sulfate-binding carbohydrate binding module of a chondroitinase (dark grey, PDB 8wab, RMSD 2.5 Å across 64 equivalent residues) which is defined as a DNRLRE domain-containing protein by UniProtKB. **c**, BamM (tan) in cartoon representation viewed from the direction indicated on the Left Hand model and overlaid with the closest structural homologue, the peptidyl-prolyl isomerases (PPI) subunit (dark grey) from the SprA Type 9 translocon complex (PDB: 6h3i chain B, RMSD 0.75 Å across 74 equivalent residues).

**Extended Data Fig. 4.**
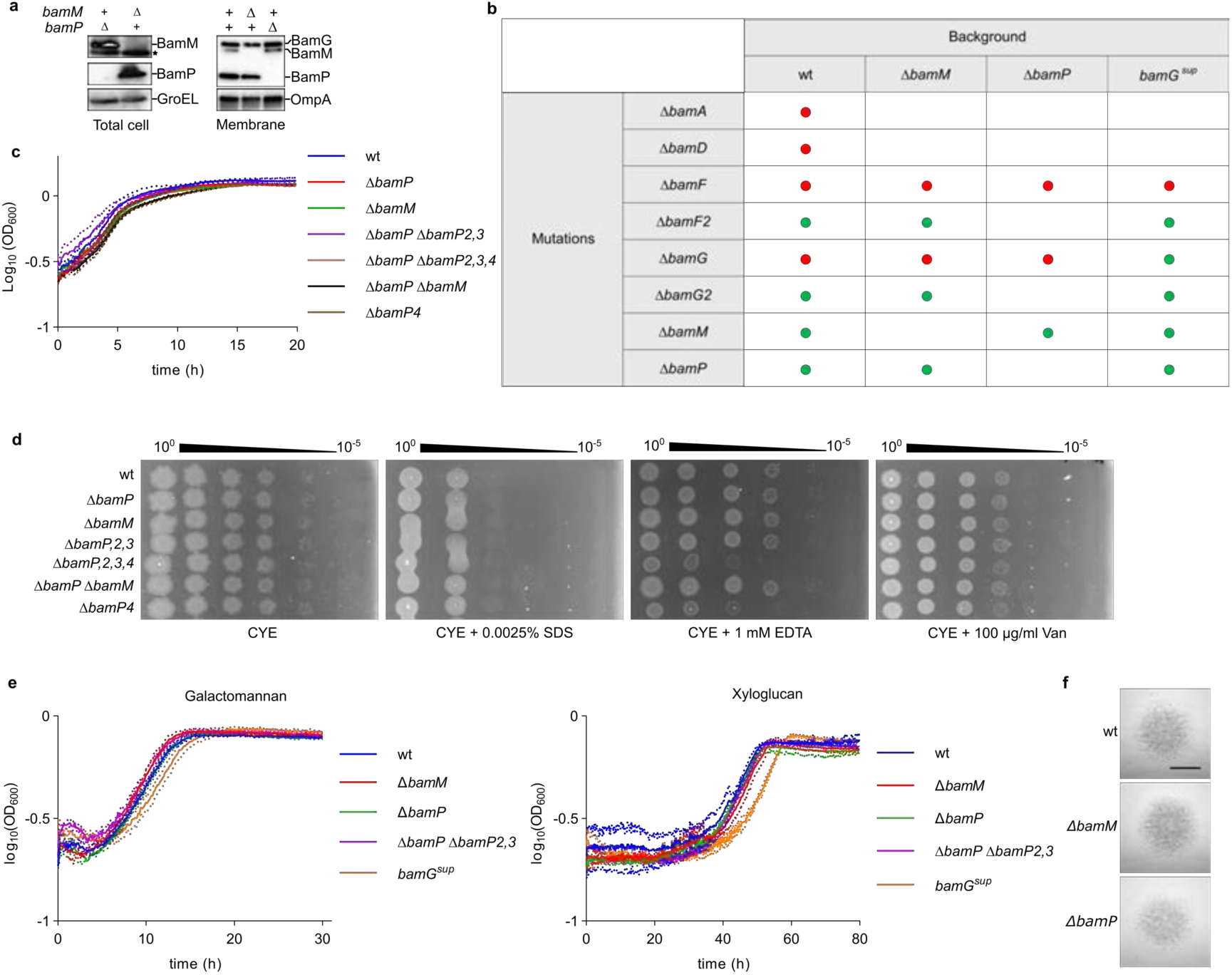
| Phenotypic characterisation of strains with deletions in BAM_Fj_ subunits or BAM_Fj_ subunit homologues. **a**, Immunoblots of whole cells and isolated membranes of strains containing in-frame deletions of *bamM* or *bamP*. The cytoplasmic protein GroEL and OM protein OmpA were used as loading control. *, non-specific band. Similar results were obtained from 3 biological repeats. **b**, Results of attempts to delete Bam_Fj_ subunits and their homologues in different genetic backgrounds. Mutant and their combinations that were viable are indicated by green dots, while those that could not be constructed are indicated by red dots and are assumed to disrupt essential cell functions. **c**, Growth curves on rich CYE medium. Shown are the means ± 1 SD from three biological repeats. **d**, OM integrity assays. Cells were grown on CYE agar with the indicated additions. Van, vancomycin. **e**, Growth curves on minimal medium containing either galactomannan or xyloglucan as carbon source. Shown are the means ± 1 SD from three biological repeats. **f**, Spreading (gliding) morphology of colonies on agar. Scale bar, 5 mm.

**Extended Data Fig. 5.**
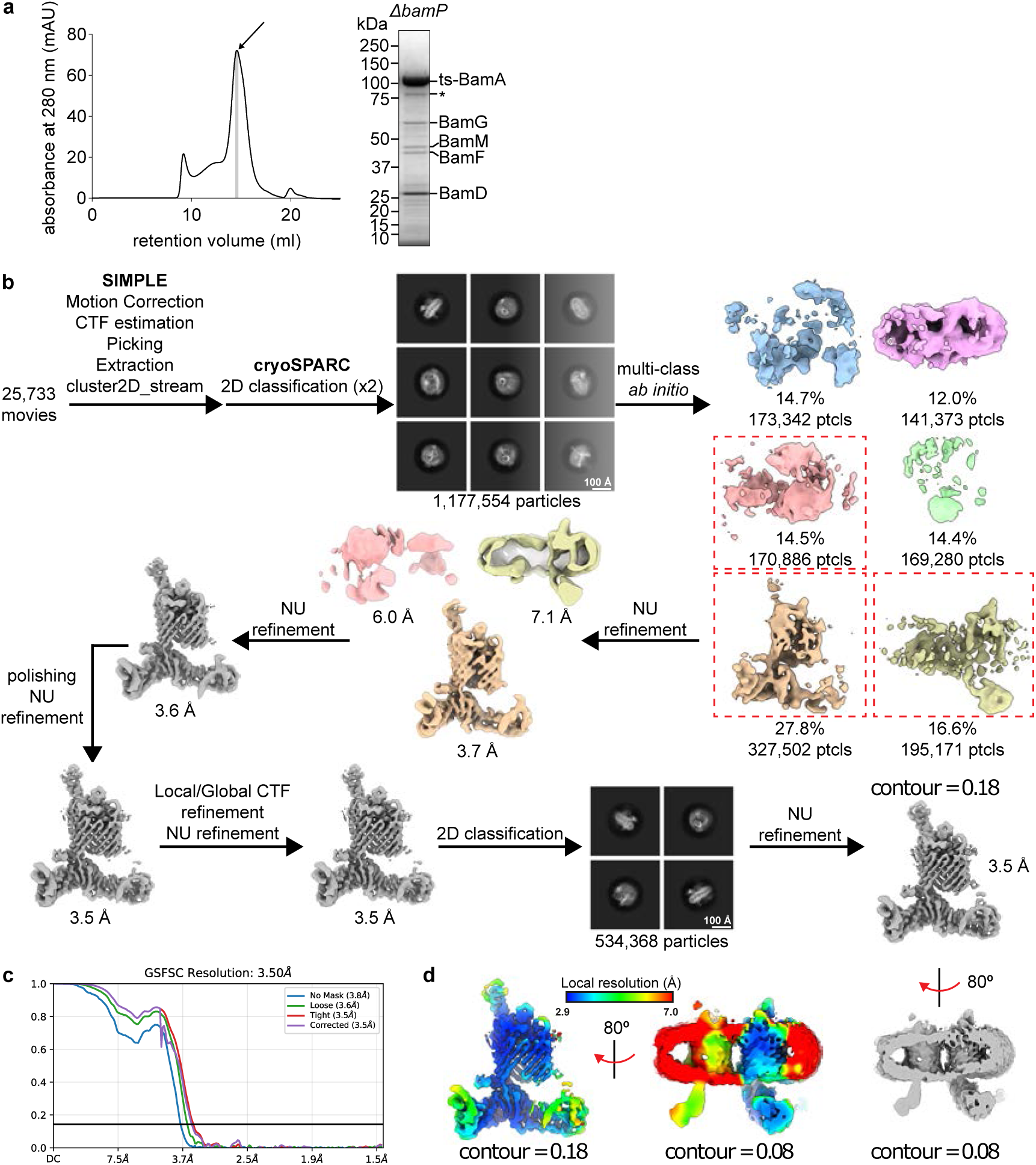
| Workflow for the cryoEM analysis of the BamA complex isolated from a Δ*bamP* background. **a**, Size exclusion chromatography profile of BamA complexes purified from a BamP-deleted background together with a Coomassie-stained SDS–PAGE gel of the indicated peak fraction that was used for structure determination. BamA* indicates a proteolysis product of BamA. Similar results were obtained from 2 biological repeats. **b**, Image processing workflow for the ΔBamP complex. **c**, Gold-standard Fourier Shell Correlation (FSC) curves used for global resolution estimation. **d**, Local resolution estimate of the volume, displayed at two contour levels.

**Extended Data Fig. 6.**
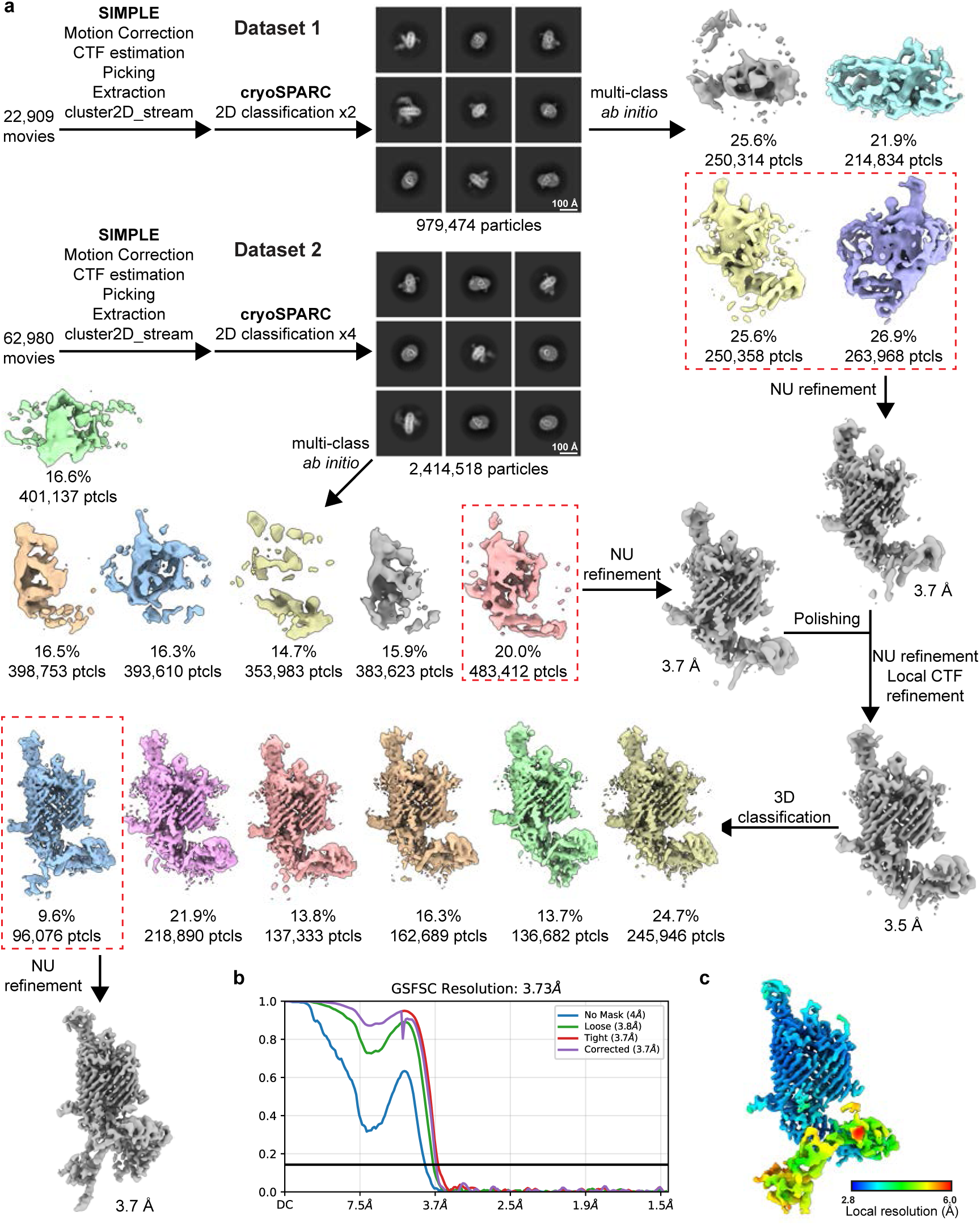
| Workflow for the cryoEM analysis of the BamAP complex. The sample used was fraction 2 from Extended Data Fig. 2a. **a**, Image processing workflow for the BamAP complexes. **b**, Gold-standard Fourier Shell Correlation (FSC) curves used for global resolution estimation. **c**, Local resolution estimate of the volume.

**Extended Data Fig. 7.**
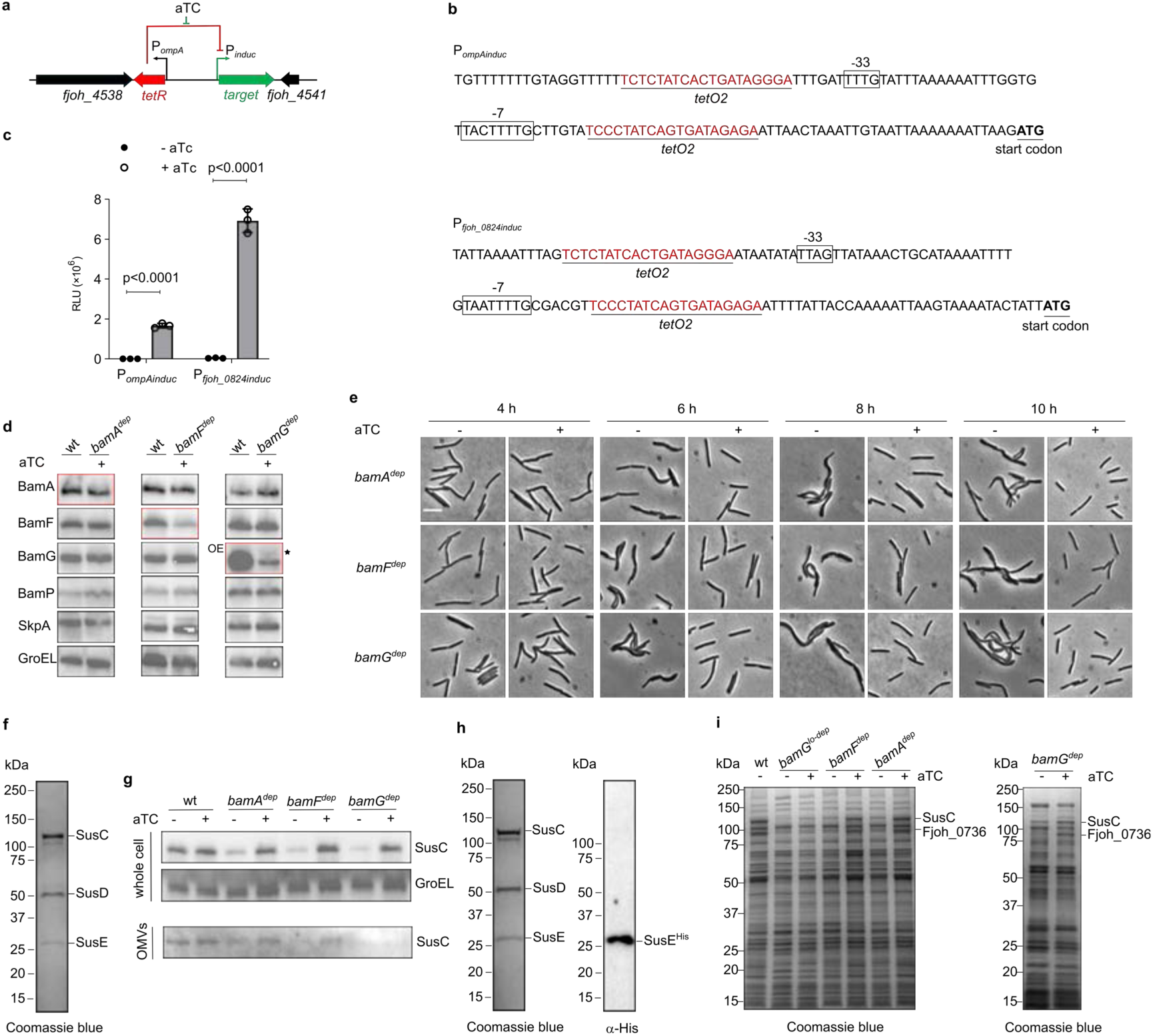
| Depletion analysis of the essential *F. johnsoniae* Bam complex subunits. **a**, Design of an anhydrotetracycline (aTC)-inducible system for the depletion of essential target genes in *F. johnsoniae*. The TetR repressor is constitutively expressed under the control of the *F. johnsoniae ompA* promoter (P*_ompA_*) and the target gene is regulated by a designed TetR-repressed promoter (P*_induc_*). In the presence of the inducer aTC repression of the target gene by TetR will be released. The genetic system is integrated into the *F. johnsoniae* chromosome at a neutral locus. **b**, Sequences of the designed inducible P*_ompA-induc_* and P*_fjoh_0824-induc_* promoters. *tetO2* arrays are placed upstream and downstream of the conserved –33 and –7 RNA polymerase binding sites (boxed) of the selected promoters. **c**, Tight regulation of protein expression by the designed inducible systems. Strains expressing NanoLuc under the control of either the P*_ompAindc_* promoter (XLFJ_1095) or the P*_fjoh_0824induc_* promoter (XLFJ_1100) were grown to mid-exponential phase (OD_600_=0.6) in the presence or absence of aTC and the luminescence signal measured. Error bars represent the mean ± 1 SD from three biological repeats. *P* values were determined with Student’s t-test. RLU, relative luminescence units. **d**, Comparison of the expression levels of Bam subunits in the wild type strain (wt, XLFJ_1078) and corresponding depletion strains grown in the presence of the inducer aTc (*bamA^dep^*, XLFJ_1129; *bamF^dep^*, XLFJ_1115; *bamG^dep^*, XLFJ_1140). Whole cell immunoblotting of cells grown to mid-exponential phase (OD_600_=0.6). The blots for the depleted subunit are boxed in red. The BamG blot for the BamG depletion comparison is overexposed (OE) relative to the other BamG blots in order to detect the low levels of BamG in the depletion strain. BamA and BamF are detected via epitope tags. *, non-specific band. **e**, Phase contrast images of cells sampled at the indicated time points in the BAM_Fj_ subunit depletion experiments shown in Fig. 4a. Scale bar, 10 µm. Similar results were obtained for 3 biological repeats. **f**, The major *F. johnsoniae* SUS complex is composed of SusC (Fjoh_0403), SusD (Fjoh_0404), and SusE (Fjoh_0405). The native SUS complex was purified via a Twin-strep tag on the N-terminus of SusC followed by size exclusion chromatography and analysed on a Coomassie-stained SDS–PAGE gel. Proteins were identified by peptide mass fingerprinting. Similar data were obtained for the two biological repeats. **g**, Outer membrane vesicle (OMV) production does not increase upon BAM depletion. Immunoblotting of the OM protein SusC in whole cells or the OMV fraction at the 6 h time point in Fig. 4a. GroEL serves as loading control. Similar results were obtained for 3 biological repeats. **h**, An exogenously-expressed His-tagged variant of SusE (SusE^His^) is incorporated into the native SusCDE complex. SusC-containing complexes were purified as described in **f** from cells expressing SusE^His^ from a plasmid. The purified material was separated by SDS-PAGE and characterized by Coomassie-staining (Left) and anti-His tag immunoblotting (Right). Similar data were obtained for two biological repeats. **i**, Exemplar Coomassie-stained SDS-PAGE gel of the whole membrane samples used for the comparative proteome analysis (Fig. 4e,g) of induced/non-induced (i.e. undepleted/depleted) BAM subunit depletion strains harvested at the 6 h time point in Fig. 4a. Proteins present in the two obviously depleting bands were assigned by peptide mass fingerprinting. The data are representative of the three repeats used for the proteomics analysis.

**Extended Data Fig. 8.**
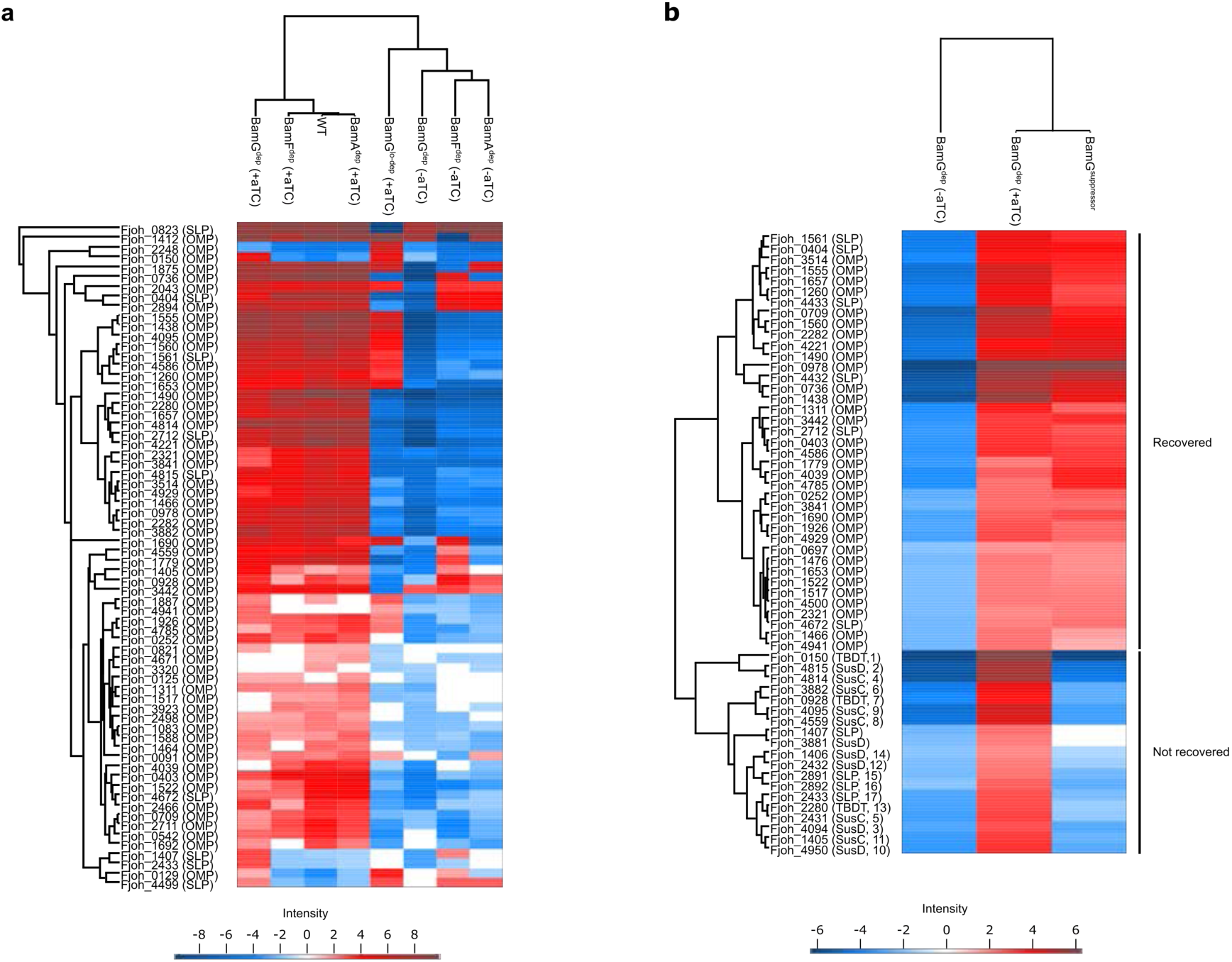
| OM proteomics data comparisons. Heat maps of the indicated strains after hierarchical protein clustering of the entire datasets and post hoc ANOVA testing. Only proteins classified as OMPs or SLPs are displayed. Colours indicate HSD (honestly significant difference) values according to the intensity panel. **a**, Comparison of the datasets used in Fig. 4e and 4g. **b**, Comparison of the *bamG^sup^* mutant dataset with the induced and non-induced *bamG^dep^* datasets. The non-recovered proteins are numbered as in Fig. 5d and assigned to SusC, SusD, other SUS SLP, or TonB-dependent transporter (TBDT) protein families. TBDTs are 22-strand OMPs that are related to the SusC family.

**Extended Data Fig. 9.**
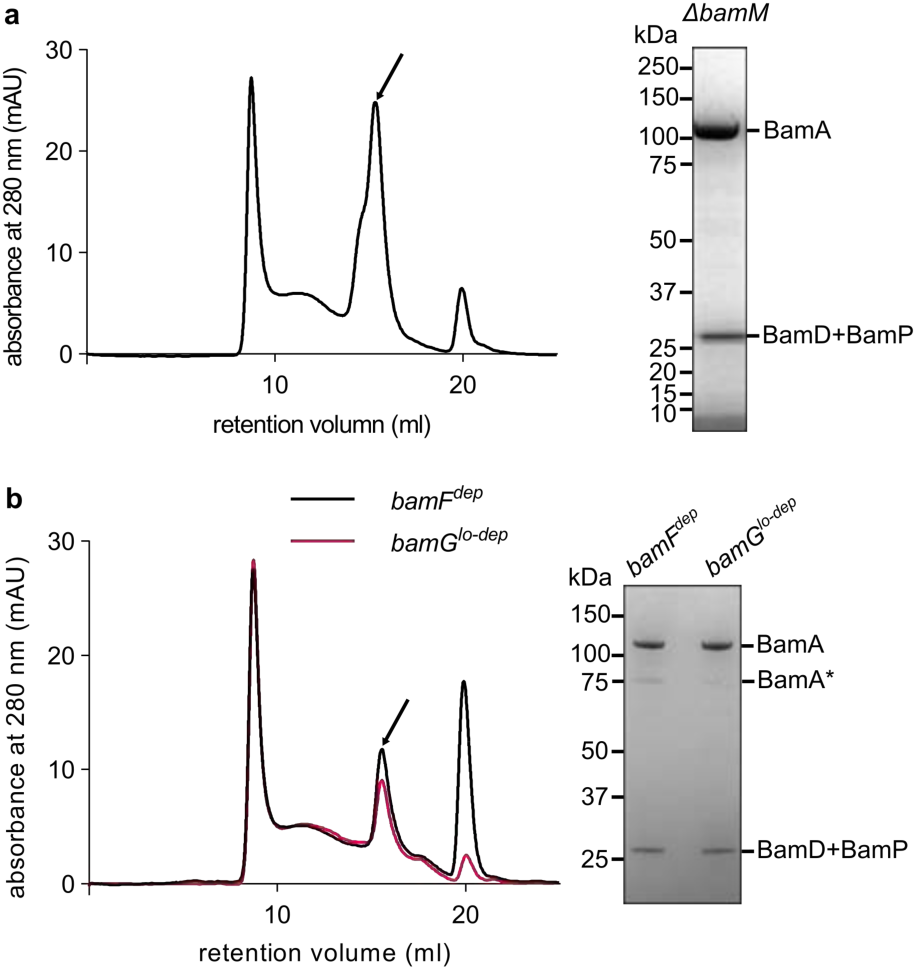
| Isolation of BamA complexes after 6 h of depletion of the essential BamF or BamG subunits or in the absence of BamM. Size exclusion chromatography profile of twin strep-tagged BamA complexes purified by streptactin affinity chromatography (Left) and a Coomassie-stained SDS–PAGE gel of the indicated peak fractions (Right). BamA* indicates a proteolysis product of BamA. The identities of the BamA* and BamD + BamP bands were assigned by peptide fingerprinting. Similar results were obtained from 2 biological repeats. **a**, Purification from the BamM deletion mutant XLFJ_998. **b**, Purifications after the depletion of BamF (strain XLFJ_1115) or BamG (strain XLFJ_1140) for 6 h.

**Extended Data Fig. 10.**
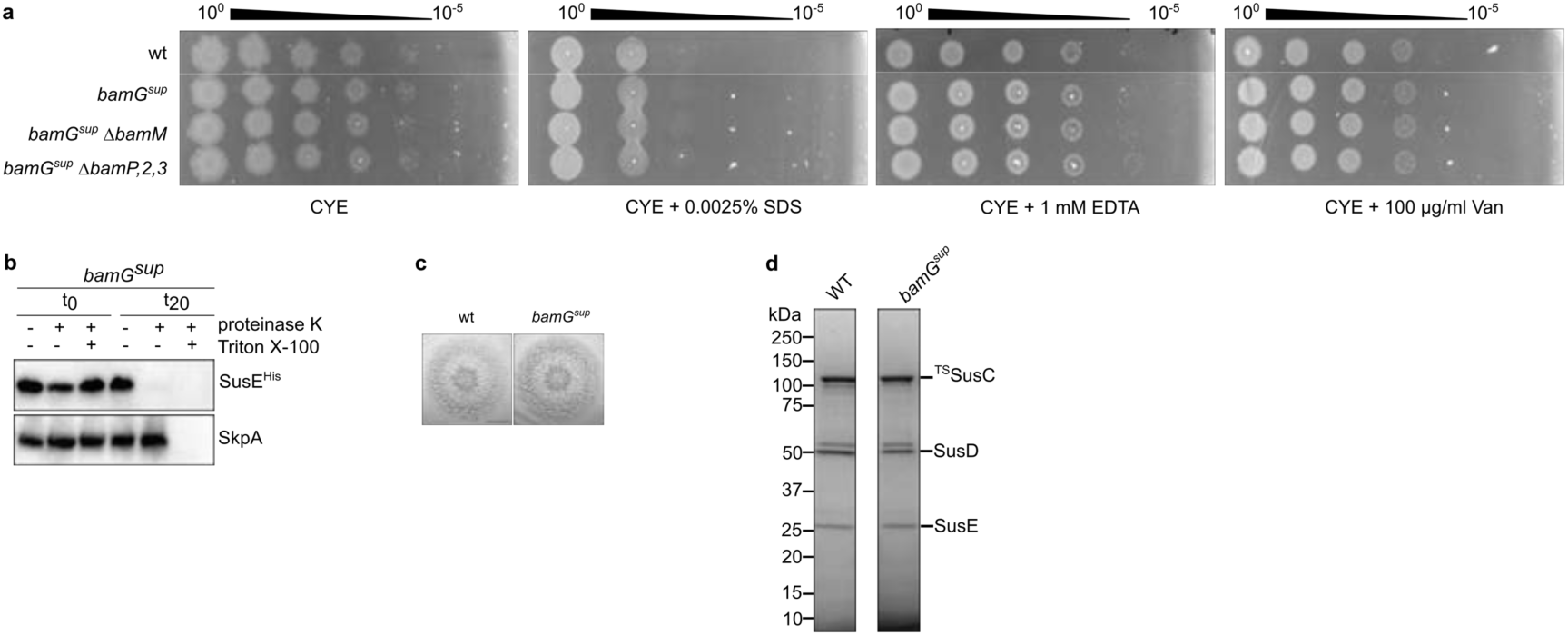
| Phenotypic characterisation of the *bamG^sup^* strain. Characterization of the recreated *bamG^sup^* mutant (*bamA^Q801K^* Δ*bamG* Δ*bamG2*). wt, wild type. Similar results were obtained for three biological repeats. **a**, OM integrity assays. Cells were grown on CYE agar with the indicated additions. Van, vancomycin. **b**, Surface exposure of the SLP SusE. Strains expressing a protease-sensitive His-tagged variant of SusE (SusE^His^) were treated as indicated with Proteinase K and the detergent Triton X-100 (to permeabilise the OM). Reactions were stopped immediately (t_0_) or after 20 min (t_20_) and analysed by immunoblotting with His tag antibodies. The periplasmic protein SkpA serves as an OM integrity control. **c**, Spreading (gliding) morphology of colonies on agar. Scale bar, 5 mm. **d**, Purification of the native SusCDE complex via a Twin-strep tag on the N-terminus of SusC followed by size exclusion chromatography. Analysed on a Coomassie-stained SDS–PAGE gel.

## Notes

### Competing Interest Statement

The authors have declared no competing interest.

## References

1 Sun, J., Rutherford, S. T., Silhavy, T. J. & Huang, K. C. Physical properties of the bacterial outer membrane. Nat Rev Microbiol 20, 236–248 (2022). 10.1038/s41579-021-00638-0

2 Konovalova, A., Kahne, D. E. & Silhavy, T. J. Outer Membrane Biogenesis. Annual review of microbiology 71, 539–556 (2017). 10.1146/annurev-micro-090816-093754

3 Combs, A. N. & Silhavy, T. J. Periplasmic Chaperones: Outer Membrane Biogenesis and Envelope Stress. Annual review of microbiology (2024). 10.1146/annurev-micro-041522-102901

4 Grabowicz, M. Lipoproteins and Their Trafficking to the Outer Membrane. EcoSal Plus 8 (2019). 10.1128/ecosalplus.ESP-0038-2018

5 Kaur, H. et al. The antibiotic darobactin mimics a beta-strand to inhibit outer membrane insertase. Nature 593, 125–129 (2021). 10.1038/s41586-021-03455-w

6 Pahil, K. S. et al. A new antibiotic traps lipopolysaccharide in its intermembrane transporter. Nature 625, 572–577 (2024). 10.1038/s41586-023-06799-7

7 Wu, T. et al. Identification of a multicomponent complex required for outer membrane biogenesis in Escherichia coli. Cell 121, 235–245 (2005). 10.1016/j.cell.2005.02.015

8 Doyle, M. T. & Bernstein, H. D. Function of the Omp85 Superfamily of Outer Membrane Protein Assembly Factors and Polypeptide Transporters. Annual review of microbiology 76, 259–279 (2022). 10.1146/annurev-micro-033021-023719

9 Takeda, H. et al. Mitochondrial sorting and assembly machinery operates by beta-barrel switching. Nature 590, 163–169 (2021). 10.1038/s41586-020-03113-7

10 Gu, Y. et al. Structural basis of outer membrane protein insertion by the BAM complex. Nature 531, 64–69 (2016). 10.1038/nature17199

11 Noinaj, N. et al. Structural insight into the biogenesis of beta-barrel membrane proteins. Nature 501, 385–390 (2013). 10.1038/nature12521

12 Bakelar, J., Buchanan, S. K. & Noinaj, N. The structure of the beta-barrel assembly machinery complex. Science 351, 180–186 (2016). 10.1126/science.aad3460

13 Tomasek, D. et al. Structure of a nascent membrane protein as it folds on the BAM complex. Nature 583, 473–478 (2020). 10.1038/s41586-020-2370-1

14 Doyle, M. T. et al. Cryo-EM structures reveal multiple stages of bacterial outer membrane protein folding. Cell 185, 1143–1156 e1113 (2022). 10.1016/j.cell.2022.02.016

15 Hartojo, A. & Doyle, M. T. beta-barrel membrane proteins fold via hybrid-barrel intermediate states. Curr Opin Struct Biol 87, 102830 (2024). 10.1016/j.sbi.2024.102830

16 Tomasek, D. & Kahne, D. The assembly of beta-barrel outer membrane proteins. Curr Opin Microbiol 60, 16–23 (2021). 10.1016/j.mib.2021.01.009

17 Human Microbiome Project, C. Structure, function and diversity of the healthy human microbiome. Nature 486, 207–214 (2012). 10.1038/nature11234

18 Silale, A. & van den Berg, B. TonB-Dependent Transport Across the Bacterial Outer Membrane. Annual review of microbiology 77, 67–88 (2023). 10.1146/annurev-micro-032421-111116

19 Horne, J. E., Brockwell, D. J. & Radford, S. E. Role of the lipid bilayer in outer membrane protein folding in Gram-negative bacteria. J Biol Chem 295, 10340–10367 (2020). 10.1074/jbc.REV120.011473

20 Lauber, F., Deme, J. C., Lea, S. M. & Berks, B. C. Type 9 secretion system structures reveal a new protein transport mechanism. Nature 564, 77–82 (2018). 10.1038/s41586-018-0693-y

21 Hooda, Y. & Moraes, T. F. Translocation of lipoproteins to the surface of gram negative bacteria. Curr Opin Struct Biol 51, 73–79 (2018). 10.1016/j.sbi.2018.03.006

22 White, J. B. R. et al. Outer membrane utilisomes mediate glycan uptake in gut Bacteroidetes. Nature 618, 583–589 (2023). 10.1038/s41586-023-06146-w

23 Veith, P. D., Glew, M. D., Gorasia, D. G., Cascales, E. & Reynolds, E. C. The Type IX Secretion System and Its Role in Bacterial Function and Pathogenesis. J Dent Res 101, 374–383 (2022). 10.1177/00220345211051599

24 Lauber, F. et al. Structural insights into the mechanism of protein transport by the Type 9 Secretion System translocon. Nat Microbiol (2024). 10.1038/s41564-024-01644-7

25 Jumper, J. et al. Highly accurate protein structure prediction with AlphaFold. Nature 596, 583–589 (2021). 10.1038/s41586-021-03819-2

26 Leonard-Rivera, M. & Misra, R. Conserved residues of the putative L6 loop of Escherichia coli BamA play a critical role in the assembly of beta-barrel outer membrane proteins, including that of BamA itself. J Bacteriol 194, 4662–4668 (2012). 10.1128/JB.00825-12

27 Heinz, E. & Lithgow, T. A comprehensive analysis of the Omp85/TpsB protein superfamily structural diversity, taxonomic occurrence, and evolution. Front Microbiol 5, 370 (2014). 10.3389/fmicb.2014.00370

28 Abramson, J. et al. Accurate structure prediction of biomolecular interactions with AlphaFold 3. Nature 630, 493–500 (2024). 10.1038/s41586-024-07487-w

29 Imai, Y. et al. A new antibiotic selectively kills Gram-negative pathogens. Nature 576, 459–464 (2019). 10.1038/s41586-019-1791-1

30 van den Berg, B., Black, P. N., Clemons, W. M., Jr. & Rapoport, T. A. Crystal structure of the long-chain fatty acid transporter FadL. Science 304, 1506–1509 (2004). 10.1126/science.1097524

31 Hearn, E. M., Patel, D. R., Lepore, B. W., Indic, M. & van den Berg, B. Transmembrane passage of hydrophobic compounds through a protein channel wall. Nature 458, 367–370 (2009). 10.1038/nature07678

32 Baez, W. D. et al. Global analysis of protein synthesis in Flavobacterium johnsoniae reveals the use of Kozak-like sequences in diverse bacteria. Nucleic Acids Res 47, 10477–10488 (2019). 10.1093/nar/gkz855

33 Hannah Martin, L. A. R., Laila Moushtaq, Amanda A. Brindley, Polly Forbes, Amy R. Quintion, Andrew R.J. Murphy, Tim J. Daniell, Didier Ndeh, Sam Amsbury, Andrew Hitchcock, Ian D.E.A. Lidbury. Hybrid xyloglucan utilisation loci are prevalent among plant-associated Bacteroidota. bioRxiv (2024). 10.1101/2024.06.03.597110

34 McBride, M. J. Bacteroidetes Gliding Motility and the Type IX Secretion System. Microbiol Spectr **7** (2019). 10.1128/microbiolspec.PSIB-0002-2018

35 Kaplan, M. et al. In situ imaging of bacterial outer membrane projections and associated protein complexes using electron cryo-tomography. Elife 10 (2021). 10.7554/eLife.73099

36 Dunstan, R. A. et al. Assembly of the secretion pores GspD, Wza and CsgG into bacterial outer membranes does not require the Omp85 proteins BamA or TamA. Mol Microbiol 97, 616–629 (2015). 10.1111/mmi.13055

37 Shibata, S. et al. Filamentous structures in the cell envelope are associated with bacteroidetes gliding machinery. Commun Biol 6, 94 (2023). 10.1038/s42003-023-04472-3

38 Lidbury, I. et al. Niche-adaptation in plant-associated Bacteroidetes favours specialisation in organic phosphorus mineralisation. The ISME journal 15, 1040–1055 (2021). 10.1038/s41396-020-00829-2

39 Kulkarni, S. S., Johnston, J. J., Zhu, Y., Hying, Z. T. & McBride, M. J. The Carboxy-Terminal Region of Flavobacterium johnsoniae SprB Facilitates Its Secretion by the Type IX Secretion System and Propulsion by the Gliding Motility Machinery. J Bacteriol 201 (2019). 10.1128/JB.00218-19

40 Wang, X., Nyenhuis, S. B. & Bernstein, H. D. The translocation assembly module (TAM) catalyzes the assembly of bacterial outer membrane proteins in vitro. Nat Commun 15, 7246 (2024). 10.1038/s41467-024-51628-8

41 Mikheyeva, I. V., Sun, J., Huang, K. C. & Silhavy, T. J. Mechanism of outer membrane destabilization by global reduction of protein content. Nat Commun 14, 5715 (2023). 10.1038/s41467-023-40396-6

42 Hart, E. M., Gupta, M., Wuhr, M. & Silhavy, T. J. The gain-of-function allele bamA(E470K) bypasses the essential requirement for BamD in beta-barrel outer membrane protein assembly. Proc Natl Acad Sci U S A 117, 18737–18743 (2020). 10.1073/pnas.2007696117

43 Takeda, H. et al. A multipoint guidance mechanism for beta-barrel folding on the SAM complex. Nat Struct Mol Biol 30, 176–187 (2023). 10.1038/s41594-022-00897-2

44 Benn, G. et al. Phase separation in the outer membrane of Escherichia coli. Proc Natl Acad Sci U S A 118 (2021). 10.1073/pnas.2112237118

45 McBride, M. J. & Kempf, M. J. Development of techniques for the genetic manipulation of the gliding bacterium Cytophaga johnsonae. J Bacteriol 178, 583–590 (1996).

46 Agarwal, S., Hunnicutt, D. W. & McBride, M. J. Cloning and characterization of the Flavobacterium johnsoniae (Cytophaga johnsonae) gliding motility gene, gldA. Proc Natl Acad Sci U S A 94, 12139–12144 (1997).

47 Gibson, D. G. et al. Enzymatic assembly of DNA molecules up to several hundred kilobases. Nat Methods 6, 343–345 (2009). 10.1038/nmeth.1318

48 Hennell James, R., et al. Structure and mechanism of the proton-driven motor that powers type 9 secretion and gliding motility. Nat Microbiol 6, 221–233 (2021). 10.1038/s41564-020-00823-6

49 Lim, B., Zimmermann, M., Barry, N. A. & Goodman, A. L. Engineered Regulatory Systems Modulate Gene Expression of Human Commensals in the Gut. Cell 169, 547–558 e515 (2017). 10.1016/j.cell.2017.03.045

50 Hall, M. P. et al. Engineered luciferase reporter from a deep sea shrimp utilizing a novel imidazopyrazinone substrate. ACS Chem Biol 7, 1848–1857 (2012). 10.1021/cb3002478

51 Cox, J. & Mann, M. MaxQuant enables high peptide identification rates, individualized p.p.b.-range mass accuracies and proteome-wide protein quantification. Nat Biotechnol 26, 1367–1372 (2008). 10.1038/nbt.1511

52 Tyanova, S. et al. The Perseus computational platform for comprehensive analysis of (prote)omics data. Nat Methods 13, 731–740 (2016). 10.1038/nmeth.3901

53 Behdenna, A. et al. pyComBat, a Python tool for batch effects correction in high-throughput molecular data using empirical Bayes methods. BMC bioinformatics 24, 459 (2023). 10.1186/s12859-023-05578-5

54 Rudolph, J. D. & Cox, J. A Network Module for the Perseus Software for Computational Proteomics Facilitates Proteome Interaction Graph Analysis. J Proteome Res 18, 2052–2064 (2019). 10.1021/acs.jproteome.8b00927

55 Teufel, F. et al. SignalP 6.0 predicts all five types of signal peptides using protein language models. Nat Biotechnol 40, 1023–1025 (2022). 10.1038/s41587-021-01156-3

56 Caesar, J. et al. SIMPLE 3.0. Stream single-particle cryo-EM analysis in real time. J Struct Biol X 4, 100040 (2020). 10.1016/j.yjsbx.2020.100040

57 Punjani, A., Zhang, H. & Fleet, D. J. Non-uniform refinement: adaptive regularization improves single-particle cryo-EM reconstruction. Nat Methods 17, 1214–1221 (2020). 10.1038/s41592-020-00990-8

58 Zivanov, J., Nakane, T. & Scheres, S. H. W. A Bayesian approach to beam-induced motion correction in cryo-EM single-particle analysis. IUCrJ 6, 5–17 (2019). 10.1107/S205225251801463X

59. UCSF pyem v0.5 (2019).

60 Pettersen, E. F. et al. UCSF ChimeraX: Structure visualization for researchers, educators, and developers. Protein Sci 30, 70–82 (2021). 10.1002/pro.3943

61 Brown, A. et al. Tools for macromolecular model building and refinement into electron cryo-microscopy reconstructions. Acta Crystallogr D Biol Crystallogr 71, 136–153 (2015). 10.1107/S1399004714021683

62 Afonine, P. V. et al. Real-space refinement in PHENIX for cryo-EM and crystallography. Acta Crystallogr D Struct Biol 74, 531–544 (2018). 10.1107/S2059798318006551

63 Williams, C. J. et al. MolProbity: More and better reference data for improved all-atom structure validation. Protein Sci 27, 293–315 (2018). 10.1002/pro.3330

